# Probiotic neoantigen delivery vectors for precision cancer immunotherapy

**DOI:** 10.1101/2023.09.29.560228

**Authors:** Andrew Redenti, Jongwon Im, Benjamin Redenti, Fangda Li, Mathieu Rouanne, Zeren Sheng, William Sun, Candice R. Gurbatri, Shunyu Huang, Meghna Komaranchath, YoungUk Jang, Jaeseung Hahn, Edward R. Ballister, Rosa L. Vincent, Ana Vardoshivilli, Tal Danino, Nicholas Arpaia

**Affiliations:** Department of Microbiology & Immunology, Vagelos College of Physicians and Surgeons of Columbia University, New York, NY 10032, USA; Department of Biomedical Engineering, Columbia University, New York, NY 10027, USA; Herbert Irving Comprehensive Cancer Center, Columbia University, New York, NY 10032, USA; Data Science Institute, Columbia University, New York, NY 10027, USA

## Abstract

Microbial systems have been synthetically engineered to deploy therapeutic payloads *in vivo*^1–4^. With emerging evidence that bacteria naturally home to tumors^5–7^ and modulate anti-tumor immunity^8,9^, one promising application is the development of bacterial vectors as precision cancer vaccines^10–12^. In this study, we engineered probiotic *E. coli* Nissle 1917 (EcN) as an anti-tumor vaccination platform optimized for enhanced production and cytosolic delivery of neoepitope-containing peptide arrays, with increased susceptibility to blood clearance and phagocytosis. These features enhance both safety and immunogenicity, achieving a system which drives potent and specific T cell–mediated anti-cancer immunity that effectively controls or eliminates tumor growth and extends survival in advanced murine primary and metastatic solid tumors. We demonstrate that the elicited anti-tumor immune response involves extensive priming and activation of neoantigen-specific CD4^+^ and CD8^+^ T cells, broader activation of both T and NK cells, and a reduction of tumor-infiltrating immunosuppressive myeloid and regulatory T and B cell populations. Taken together, this work leverages the advantages of living medicines to deliver arrays of tumor-specific neoantigen–derived epitopes within the optimal context to induce specific, effective, and durable systemic anti-tumor immunity.

## Main

Bacteria support activation of both innate and adaptive immunity through their inherent foreignness and immunostimulatory properties^13,14^. These features, coupled with the ease to synthetically engineer them for safe delivery of immunomodulatory compounds, make bacteria ideal vectors to augment and direct anti-tumor immune responses^2–4^. Tumor neoantigens are attractive immunotherapeutic payloads for delivery; these antigenic species are not present in other tissues, pose minimal risk for inducing autoimmunity, and are theoretically excluded from central immunologic tolerance mechanisms^15,16^. To date, a variety of tumor neoantigen vaccines have demonstrated promising immunologic responses and survival benefit in clinical trials, though benefit remains limited to only a subset of patients^17–20^. In this regard, programming bacteria with genetic directives to release high levels of identified tumor neoantigens thereby provides a system to precisely instruct neoantigen targeting *in situ*. Herein, we describe novel microbial immunotherapy vectors that stimulate effective and durable tumor antigen–specific immunity and inhibit immunosuppressive mechanisms that may otherwise limit traditional neoantigen vaccines^21^.

### Engineering microbes for tumor vaccination

To enable effective cancer vaccination, we developed an engineered bacterial system in probiotic EcN to enhance expression, delivery, and immune-targeting of arrays of tumor exonic mutation– derived epitopes highly expressed by tumor cells and predicted to bind major histocompatibility complex (MHC) class I and II (**Fig. 1a**). This system incorporates several key design elements that enhance therapeutic utility: *(1)* optimization of synthetic neoantigen construct form with *(2)* removal of cryptic plasmids and deletion of Lon and OmpT proteases to increase neoantigen accumulation, *(3)* increased susceptibility to phagocytosis for enhanced uptake by antigen-presenting cells (APCs) and presentation of MHC class II–restricted antigens, *(4)* expression of Listeriolysin O (LLO) to induce cytosolic entry for presentation of recombinant encoded neoantigens by MHC class I molecules and Th1-type immunity, and *(5)* improved safety for systemic administration due to reduced survival in the blood and biofilm formation.

**Figure 1.**
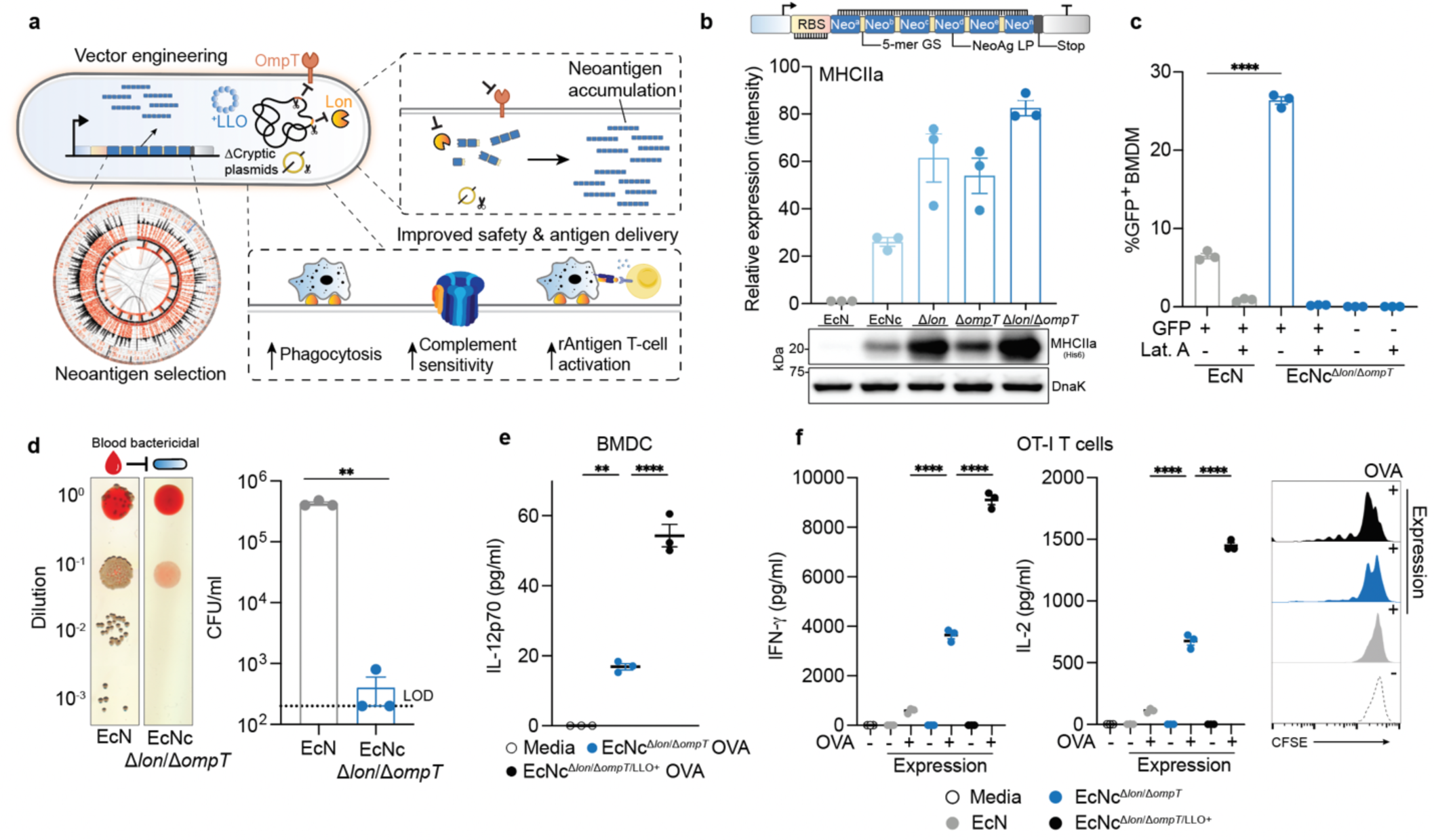
Engineering live microbial tumor neoantigen vaccines. **a**, Design of microbial tumor neoantigen vectors, with Circos plot of CT26 mutanome. **b**, Upper: optimized synthetic neoantigen construct schematic. Middle: relative immunoblot chemiluminescent intensity of neoantigen construct MHCIIa expressed from parent (EcN) vs. derivative strains (*n* = 3 replicates per group). Lower: representative immunoblot of neoantigen construct MHCIIa expression in wildtype EcN, EcNc, EcNc^Δ*lon*^, EcNc^Δ*ompT*^, or EcNc^Δ*lon*/Δ*ompT*^. **c**, Percent GFP^+^ BMDM after incubation with EcN or EcNc^Δ*lon*/Δ*ompT*^ expressing constitutive GFP (*n* = 3 replicates per group, \*\*\*\**P* < 0.0001, unpaired student’s t-test); Lat. A = Latrunculin A. **d**, Left: representative image of EcN or EcNc^Δ*lon*/Δ*ompT*^ spotted on LB agar plate after incubation in human blood, Right: microbial burden quantified as CFU/ml (*n* = 3 replicates per group, \*\**P* = 0.0039, unpaired student’s t-test with Welch’s correction). Limit of detection (LOD) = 2 × 10^2^ CFU/ml **e**, IL-12p70 quantification in culture supernatants of BMDC pulsed with indicated condition (*n* = 3 replicates per group, \*\**P* = 0.0018, \*\*\*\**P* < 0.0001, one-way ANOVA with Tukey’s multiple comparisons test). **f**, Naïve OT-I T cells were incubated with BMDC’s pulsed with the indicated condition. Left: IFN-γ quantification in supernatants of OT-I cultures (*n* = 3 replicates per group, \*\*\*\**P* < 0.0001, one-way ANOVA with Tukey’s multiple comparisons test), Middle: IL-2 quantification in supernatants of OT-I cultures (*n* = 3 replicates per group, \*\*\*\**P* < 0.0001, one-way ANOVA with Tukey’s multiple comparisons test). Right: representative histogram depicting CFSE dilution of stimulated OT-I T cells. **b**–**f,** Data are mean ± s.e.m.

To assemble a repertoire of neoantigens, we conducted exome and transcriptome sequencing of subcutaneous CT26 tumors. Neoantigens were predicted from highly expressed tumor-specific mutations using established methods^22,23^, with selection criteria inclusive of putative neoantigens across a spectrum of MHC affinity^24–26^. Given the importance of both MHC class I and MHC class II binding epitopes in anti-tumor immunity^23,27,28^, we integrated a measure of wildtype-to-mutant MHC affinity ratio – agretopicity or MHC amplitude^26,29,30^ – for both epitope types derived from a given mutation, to estimate the ability of adaptive immunity to recognize a neoantigen. Predicted neoantigens were selected from the set of tumor-specific mutations satisfying all criteria, notably encompassing numerous recovered, previously validated CT26 neoantigens^23^ (**Extended Data Fig. 1a**).

We then sought to create a microbial system which could accommodate the production and delivery of diverse sets of neoantigens to lymphoid tissue and the tumor microenvironment (TME). For the purpose of assessing neoantigen production capacity, a prototype gene encoding a synthetic neoantigen construct (NeoAg^p^) was created by concatenating long peptides (LPs) encompassing linked CD4^+^ and CD8^+^ T cell mutant epitopes – previously shown as an optimal form for stimulating cellular immunity^31–33^ – derived from CT26 neoantigens (**Extended Data Fig. 1b, Extended Data Table 1**). The construct was cloned into a stabilized plasmid under constitutive expression and transformed into EcN, however, both immunoblot and ELISA assessment revealed low production of the prototype construct by EcN across several tested promoters (**Extended Data Fig. 1c**).

Given the dependency on antigen dosage for establishing an effective and immunodominant antigen-specific immune response^34–40^, we developed a system for improved recombinant neoantigen construct production. The incorporation of 5-mer glycine-serine (GS) linkers between neoantigen LPs in the prototype increased expression approximately 6-fold (**Extended Data Fig. 1c, d**). Conversely, expressing only minimal neoepitopes, decreasing the number of neoantigen LPs in a construct, or incorporating 10-mer GS or immunoprotease-sensitive linkers did not improve production (**Extended Data Fig. 1e**). To evaluate the capacity of constructs with 5-mer GS linkers to accommodate production of various neoantigens, and for eventual *in vivo* testing, we created 3 additional constructs with unique neoantigens from the predicted set, selected on a spectrum of predicted affinity for MHC class I and MHC class II (MHCIa, MHCIIa, MHCI/II^v^) (**Extended Data Table 2**). Prototype and novel construct expression was evaluated in EcN versus BL21, a strain which harbors chromosomal deletions of the Lon (Δ*lon*) and OmpT (Δ*ompT*) proteases to facilitate recombinant protein production^41^. Unlike BL21, EcN also bears cryptic plasmids which can suppress the copy number of transformed recombinant plasmids^42^. Indeed, on average, BL21 produced 10-fold higher levels of neoantigen construct relative to EcN (**Extended Data Fig. 1f**). Thus, to further enhance construct expression in EcN, we performed sequential synthetic modifications of the microbe. Removal of the EcN cryptic plasmids led to maintenance of approximately 300-fold higher levels of therapeutic plasmid DNA (EcNc) (**Extended Data, Fig. 2a**), with successive deletion of the Lon protease (EcNc^Δ*lon*^), OmpT protease (EcNc^Δ*ompT*^), or both proteases (EcNc^Δ*lon/*Δ*ompT*^) allowing up to 80-fold increased production of synthetic neoantigen constructs relative to the parental EcN strain (**Fig. 1b, Extended Data Fig. 2b, c**).

Since the Lon protease has been connected to capsule and biofilm regulation^43,44^, and OmpT with the degradation of complement^45^, we tested the susceptibility of the engineered vector EcNc^Δ*lon/*Δ*ompT*^ to phagocytosis and blood clearance, as well as for its proficiency in biofilm formation. Notably, EcNc^Δ*lon*/Δ*ompT*^ was 4-fold more susceptible to phagocytosis by bone marrow-derived macrophages (BMDMs) relative to EcN (**Fig. 1c**). Incubation in human blood further revealed a 100-fold greater sensitivity to blood clearance for EcNc^Δ*lon*/Δ*ompT*^ vs. EcN (**Fig. 1d**). Moreover, EcNc^Δ*lon*/Δ*ompT*^ was significantly attenuated in biofilm formation, a major mechanism of microbial resistance to immunity and antimicrobial agents in humans^46^ (**Extended Data, Fig. 2d**).

As an anti-tumor vaccine, the microbial platform must additionally facilitate presentation of recombinant antigens and activation of APCs. To evaluate the system in this capacity, the model antigen ovalbumin (OVA) was expressed in the cytosol of EcNc^Δ*lon*/Δ*ompT*^ using a strategy analogous to that used for synthetic neoantigen constructs. BMDMs pulsed with EcNc^Δ*lon*/Δ*ompT*^-OVA, but not EcN-OVA, presented the H2K^b^-SIINFEKL complex, indicating efficient processing and cross-presentation of recombinant antigens from EcNc^Δ*lon*/Δ*ompT*^ (**Extended Data Fig 2e**). Furthermore, pulsed BMDMs upregulated MHC class II and CD80, and downregulated PD-L1, demonstrating effective APC activation by EcNc^Δ*lon*/Δ*ompT*^ expressing a recombinant antigen (**Extended Data Fig. 2e, f**).

To refine the immune program orchestrated by APCs, we reasoned that constitutive co-expression of LLO – a pH-dependent protein derived from *Listeria* which forms pores in the phagolysosomal membrane – would enhance cytosolic delivery of encoded neoantigens for presentation to CD8^+^ T cells, and support APC activation and induction of Th1 immunity^47–50^. Of note, engineered microbes produced functional LLO, and LLO expression did not affect viability of antigen-presenting cells incubated with LLO-expressing strains (EcNc^Δ*lon*/Δ*ompT*/LLO+^) or the co-expression of neoantigen constructs (**Extended Data Fig. 2g, h**). Indeed, bone-marrow derived dendritic cells (BMDCs) pulsed with live EcNc^Δ*lon*/Δ*ompT*/LLO+^ OVA secreted 3-fold higher levels of IL-12p70 as compared to those pulsed with EcNc^Δ*lon*/Δ*ompT*^ OVA (**Fig. 1e**), indicating greater Th1 instruction by APCs. Moreover, BMDCs pulsed with live EcNc^Δ*lon*/Δ*ompT*/LLO+^ OVA mediated superior activation of naïve OT-I and OT-II T cells, with 2-to 2.5-fold increased secretion of interferon-γ (IFN-γ) and interleukin-2 (IL-2) from both T cell types relative to EcNc^Δ*lon*/Δ*ompT*^ OVA, and marked proliferation of both OT-I and OT-II T cells (**Fig. 1f**, **Extended Data, Fig. 2i**). Conversely, BMDCs pulsed with wildtype EcN OVA induced no measurable proliferation of either T cell type, 13-to 15-fold lower secretion of IL-2 and IFN-γ from OT-I T cells, and no detectable cytokine secretion from OT-II T cells (**Fig. 1f**, **Extended Data, Fig. 2i**). Taken together, these data suggest recombinant antigens expressed in EcNc^Δ*lon*/Δ*ompT*/LLO+^ lead to potent antigen-specific activation of both naïve cytotoxic and helper T cells, with incorporation of LLO facilitating both enhanced presentation to CD8^+^ T cells and Th1-type immunity.

Overall, synthetic neoantigen construct optimization and genetic engineering achieved a microbial platform (EcNc^Δ*lon*/Δ*ompT*/LLO+^) capable of robust production across diverse sets of tumor neoantigens, which was attenuated in immune-resistance mechanisms, effectively taken up by and proficient in activating antigen-presenting cells, and able to drive potent activation of T cells specific for encoded recombinant antigens to support enhanced cellular immunity.

### Microbial efficacy in murine colorectal carcinoma

To assess the *in vivo* efficacy of the engineered system, BALB/c mice bearing advanced CT26 tumors on a single hind-flank received an intratumoral injection of EcN wildtype, EcNc^Δ*lon*/Δ*ompT*^ or EcNc^Δ*lon*/Δ*ompT*/LLO+^ strains. These strains were tested either without any neoantigen plasmid (NC), expressing a single neoantigen construct (MHCIa, MHCIIa, and MHCI/II^v^), or as a combination of the 3 neoantigen construct–expressing strains in equal parts – a microbial anti-tumor vaccine delivering 19 total unique neoantigens (nAg^19^). Notably, no difference in tumor colonization efficiency was observed for EcNc^Δ*lon*/Δ*ompT*^ strains as compared to wildtype EcN (**Extended Data Fig. 3a, b**). Whereas intratumoral treatment with wildtype EcN expressing any neoantigen construct did not demonstrate therapeutic benefit (**Extended Data Fig. 3c, d**), a single intratumoral injection of EcNc^Δ*lon*/Δ*ompT*/LLO+^ nAg^19^ provided strong anti-tumor efficacy, with a complete response observed for 3 out of 7 tumors and the combination nAg^19^ more effective than any construct alone (**Fig. 2a, Extended Data Fig. 3e–g**). Moreover, treatment with EcNc^Δ*lon*/Δ*ompT*^ and EcNc^Δ*lon*/Δ*ompT*/LLO+^ strains was well-tolerated, with significantly attenuated body weight change as compared to wildtype EcN, and no significant body weight differences as compared to PBS treatment over the course of observation (**Extended Data Fig. 4a**). Direct comparisons of intratumoral treatment with EcNc^Δ*lon*/Δ*ompT*/LLO+^ nAg^19^ vs. EcNc^Δ*lon*/Δ*ompT*^ nAg^19^ revealed that the inclusion of LLO significantly enhanced tumor control and extended survival (**Fig. 2a, Extended Data Fig. 4b–d**). Thus, the combination of all synthetic modifications (EcNc^Δ*lon*/Δ*ompT*/LLO+^ nAg^19^) synergized to produce a microbial anti-tumor vaccine with favorable toxicity profile and strong therapeutic effect *in vivo*.

**Figure 2.**
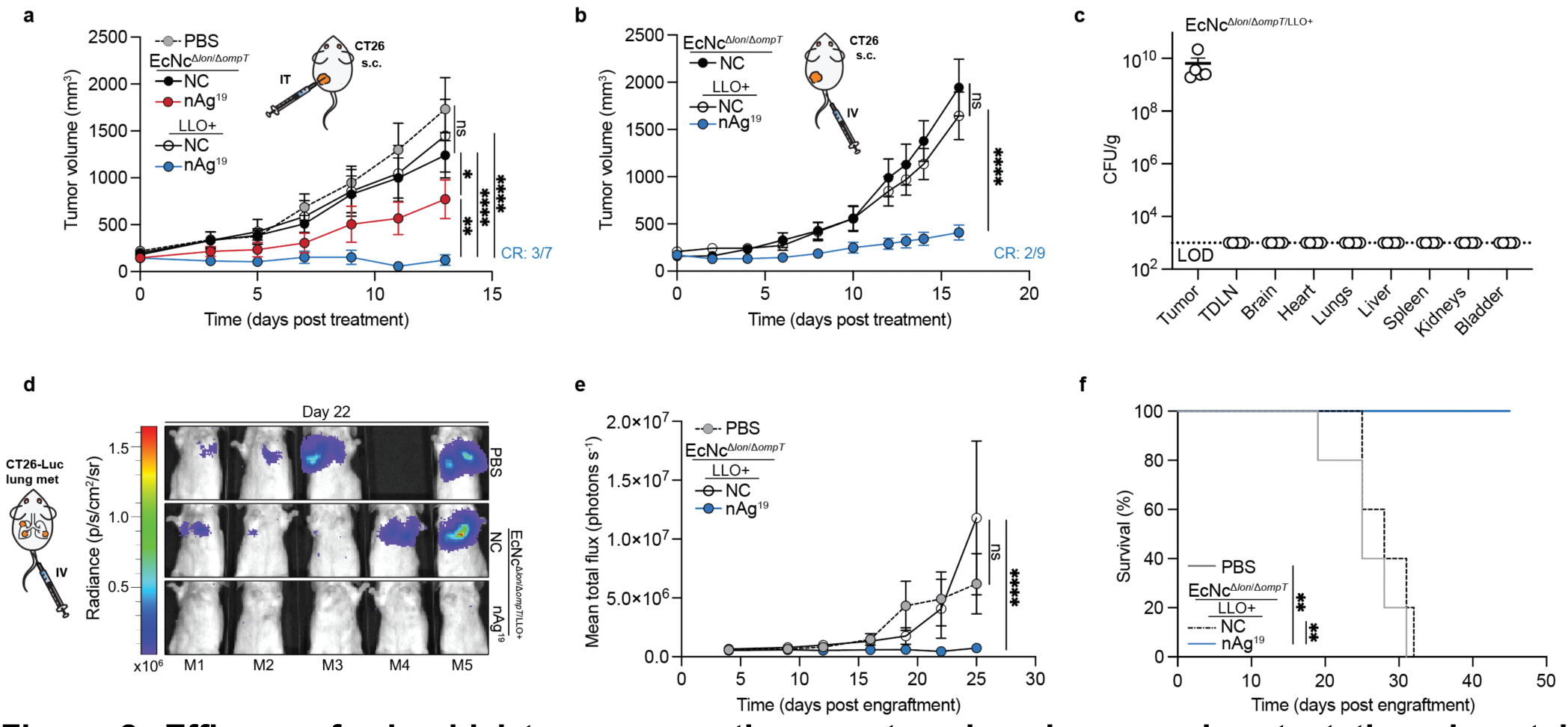
Efficacy of microbial tumor neoantigen vectors in primary and metastatic colorectal carcinoma. **a–c**, BALB/c mice with established hind-flank CT26 tumors were treated when average tumor volume was 150–200mm^3^. **a**, Mice received an intratumoral injection of PBS, EcNc^Δ*lon*/ΔompT^ or EcNc^Δ*lon*/ΔompT/LLO+^ without neoantigens (NC) or the 3 neoantigen expressing strain combination (nAg^19^) on day 0. Tumor growth curves (*n* = 5–7 mice per group, \**P* = 0.0469, \*\**P* = 0.0096, \*\*\*\**P* < 0.0001, ns = not significant (*P* > 0.05), two-way ANOVA with Tukey’s multiple comparisons test). **b**, Mice received an intravenous injection of either EcNc^Δ*lon*/Δ*ompT*^ NC, EcNc^Δ*lon*/Δ*ompT*/LLO+^ NC, or EcNc^Δ*lon*/Δ*ompT*/LLO+^ nAg^19^ on day 0 and day 8. Tumor growth curves (*n* = 8–9 mice per group, \*\*\*\**P* < 0.0001, ns = not significant (*P* > 0.05), two-way ANOVA with Tukey’s multiple comparisons test). **c**, Mice (*n* = 5) received an intravenous injection of EcNc^Δ*lon*/Δ*ompT*/LLO+^. Microbial tissue burden quantified as colony-forming units (CFU) per gram of tissue (CFU/g), LOD = 1 × 10^3^ CFU/g. **d–f**, BALB/c mice were injected intravenously with CT26-Luc cells. Starting day 4 post tumor cell injection, mice (*n* = 5 per group) received intravenous injections of PBS, EcNc^Δ*lon*/Δ*ompT*/LLO+^ NC, or EcNc^Δ*lon*/Δ*ompT*/LLO+^ nAg^19^ every 3–5 days. **d**, Representative images of lung metastases luminescence in each mouse (M1–M5) per group on day 22 post engraftment. **e**, Mean total flux from lung metastases (*n* = 5 mice per group, \*\*\*\**P* > 0.0001, ns = not significant (*P* > 0.05), two-way ANOVA with Dunnett’s multiple comparisons test). **h**, Kaplan-Meier survival curve for mice with CT26-Luc lung metastases (*n* = 5 mice per group, \*\**P* = 0.0017, \*\**P* = 0.0018, Log-Rank Mantel-Cox test). **a**–**d**, **f**, Data are mean ± s.e.m.

To evaluate the induction of systemic anti-tumor immunity after treatment with the microbial neoantigen vaccines, mice with established CT26 tumors on both hind-flanks were treated with an injection of microbes into a single tumor. Whereas treatment with EcNc^Δ*lon*/Δ*ompT*/LLO+^ without neoantigen expression (NC) did not suppress tumor growth, a single injection of EcNc^Δ*lon*/Δ*ompT*/LLO+^ nAg^19^ induced tumor control and complete regression of 2 of 6 treated and untreated tumors (**Extended Data Fig. 5a**). Microbial quantification from tumors 14-days after injection revealed that microbes colonized treated tumors at high densities, with no bacteria able to be cultured from untreated tumors (**Extended Data Fig. 5b**). This demonstrates that the engineered neoantigen vaccines stimulate systemic anti-tumor immunity capable of eliminating distant tumors.

We next evaluated the efficacy of our microbial anti-tumor vaccination platform following intravenous administration, the preferred route of administration as to circumvent dependance on tumor accessibility. Similar to intratumoral treatment, intravenous administration of EcNc^Δ*lon*/Δ*ompT*/LLO+^ nAg^19^ to mice with advanced CT26 tumors again provided potent anti-tumor efficacy and survival benefit with minimal body weight alteration (**Fig. 2b, Extended Data Fig. 6a–c**). After intravenous injection, the engineered microbes persisted at high density within tumors and were cleared rapidly from all other surveyed organs (**Fig. 2c, Extended Data Fig. 6d**).

Having observed robust efficacy via intravenous delivery, we then assessed therapeutic efficacy against established metastatic disease. CT26 carcinoma cells with genomically-integrated firefly luciferase (CT26-Luc) were injected intravenously, which form rapidly progressive lung-metastases traceable by bioluminescence imaging. Intravenous treatment with PBS, EcNc^Δ*lon*/*ΔompT*/LLO+^ without neoantigen expression (NC), or EcNc^Δ*lon*/Δ*ompT*/LLO+^ nAg^19^ was initiated 4-days after engraftment. We found that engineered microbes colonized metastases-bearing lungs and were not detectable in other tissues (**Extended Data. Fig 6e**). Microbial treated groups again demonstrated minimal body weight fluctuation, similarly to mice treated with PBS (**Extended Data Fig. 6f**). Treatment with EcNc^Δ*lon*/Δ*ompT*/LLO+^ nAg^19^ strongly restrained metastatic growth, with 100% of neoantigen therapeutic-treated mice surviving to 45 days after engraftment, whereas both control groups had completely succumbed to disease (**Fig. 2d**–**f****, Extended Data Fig. 6g**). This demonstrates both safety and efficacy of intravenously administered EcNc^Δ*lon*/Δ*ompT*/LLO+^ nAg^19^ in the setting of aggressive, established metastatic disease.

### Dynamics of microbial-stimulated immunity in colorectal carcinoma

As the engineered microbial neoantigen vaccines are strong immunostimulants and persist within the TME, we reasoned that sustained intratumoral neoantigen production and reduced immunosuppression would facilitate enhanced activation of adaptive immunity to mediate the observed tumor control. To confirm *in situ* delivery of encoded neoantigens, we intravenously administered EcNc^Δ*lon*/Δ*ompT*/LLO+^ nAg^19-His^ (wherein all three neoantigen constructs contain a C-terminal 6xHis-tag) and performed immunoblots of tumor and tumor draining lymph node (TDLN) lysates 2 days following treatment. We observed 3 His-tagged protein species corresponding to the 3 encoded neoantigen constructs (**Fig. 3a**). Consistent with delivered neoantigens enhancing T cell activation, *ex vivo* restimulation of lymphocytes isolated from TDLNs at 8 days post-treatment with phorbol myristate acetate (PMA) and ionomycin revealed increased production of IFN-γ and TNF-α by conventional CD4^+^ (Foxp3^-^CD4^+^) and CD8^+^ T cells in mice treated with EcNc^Δ*lon*/Δ*ompT*/LLO+^ nAg^19^ vs. those treated with PBS or control bacteria (EcNc^Δ*lon*/Δ*ompT*^) (**Extended Data Fig. 7a–b**).

**Figure 3.**
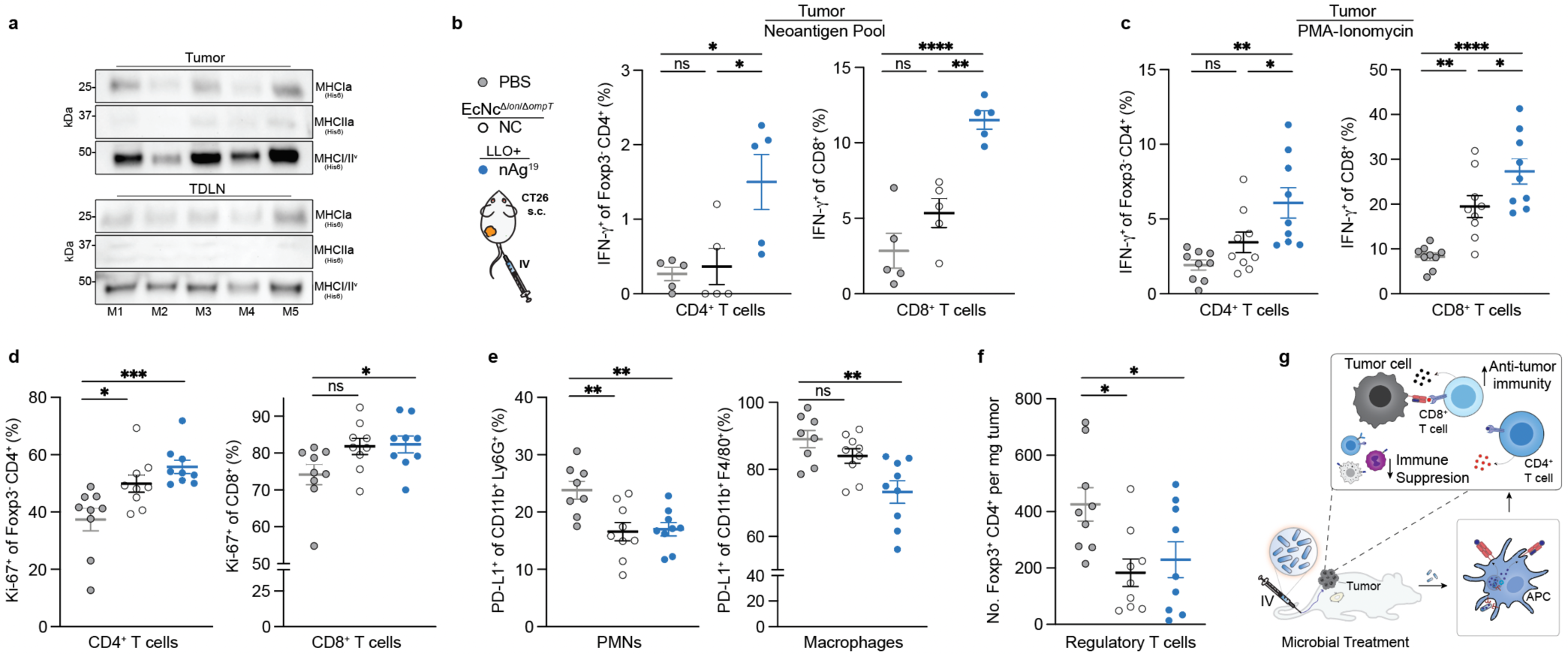
Microbial tumor neoantigen vaccines drive anti-tumor immunity and remodel the tumor immune microenvironment. **a**, Immunoblot (anti-6×His) of tumors (*n* = 5) and TDLNs (*n* = 5) from mice (M1–M5) treated intravenously with strain mixture nAg^19-His^ expressing neoantigen constructs MHCIa_(His6)_, MHCIIa_(His6)_, and MHCI/II^v^_(His6)_. **b**, Tumor-infiltrating lymphocytes were stimulated with pooled peptides representing the neoantigens encoded in EcNc^Δ*lon*/Δ*ompT*/LLO+^ nAg^19^ in the presence of brefeldin A. Left: Frequency of IFN-γ_+_ Foxp3^-^CD4^+^ post-stimulation (*n* = 5 mice per group, \**P* = 0.0238, \**P* = 0.0147, ns = not significant (*P* > 0.05), one-way ANOVA with Tukey’s multiple comparisons test). Right: Frequency of IFN-γ_+_ CD8^+^ T cells post-stimulation (*n* = 5 mice per group, \*\**P* = 0.0015, \*\*\*\**P* < 0.0001, ns = not significant (*P* > 0.05), one-way ANOVA with Tukey’s multiple comparisons test). **c**, Tumor-infiltrating lymphocytes were stimulated with PMA and ionomycin in the presence of brefeldin A. Left: Frequency of IFN-γ_+_ Foxp3^-^CD4^+^ post-stimulation (*n* = 9 mice per group, \**P* = 0.0461, \*\**P* = 0.0014, ns = not significant (*P* > 0.05), one-way ANOVA with Tukey’s multiple comparisons test). Right: frequency of IFN-γ_+_ CD8^+^ T cells post-stimulation (*n* = 9 mice per group, \**P* = 0.0486, \*\**P* = 0.0040, \*\*\*\**P* < 0.0001, one-way ANOVA with Tukey’s multiple comparisons test). **d**, Left: Percentage Ki-67^+^ of Foxp3^-^CD4^+^ T cells in tumors (*n* = 9 mice per group, \**P* = 0.0183, \*\*\**P* = 0.0008, one-way ANOVA with Dunnett’s multiple comparisons test). Right: Percentage Ki-67^+^ of CD8^+^ T cells in tumors (*n* = 9 mice per group, \**P* = 0.0453, ns = not significant (*P* > 0.05), One-way ANOVA with Dunnett’s multiple comparisons test). **e**, Left: Percentage PD-L1^+^ of Ly6g^+^CD11b^+^ in tumors (*n* = 8-9 mice per group, \*\**P* = 0.0037, \*\**P* = 0.0059, one-way ANOVA with Dunnett’s multiple comparisons test). Right: Percentage PD-L1^+^ of CD11b^+^F4/80^+^ in tumors (*n* = 8-9 mice per group, \*\**P* = 0.0010, ns = not significant (*P* > 0.05), One-way ANOVA with Dunnett’s multiple comparisons test). **f**, Number of Foxp3^+^CD4^+^ T cells per mg tumor (*n* = 9 mice per group \**P* = 0.0131, \**P* = 0.0241, one-way ANOVA with Holm-Šídák’s multiple comparisons test). **g**, Immunologic mechanism. **b**–**f**, Data are mean ± s.e.m.

Next, to assess the ability for engineered neoantigen therapeutics to drive neoantigen-specific immunity, tumor-infiltrating lymphocytes (TILs) were isolated at 8 days post-treatment and restimulated *ex vivo* with a pool of synthetic peptides representing the 19 bacterially-encoded tumor neoantigens. Flow cytometric analysis revealed increased frequencies of IFN-γ secreting conventional Foxp3^-^CD4^+^ and CD8^+^ TILs, demonstrating that treatment with EcNc^Δ*lon*/Δ*ompT*/LLO+^ nAg^19^ enhanced encoded neoantigen-specific immunity (**Fig. 3b**). Compared to peptide stimulation, restimulation with PMA and ionomycin revealed even greater levels of IFN-γ secreting Foxp3^-^CD4^+^ and CD8^+^ TILs in EcNc^Δ*lon*/Δ*ompT*/LLO+^ nAg^19^–treated tumors, and IFN-γ producing B220^+^ B-cells^51,52^, suggestive of epitope spreading and expanded immune activation (**Fig. 3c, Extended Data Fig. 7c**)^53^. Additionally, we observed increased frequencies of proliferating CD4^+^ and CD8^+^ tumor-infiltrating T cells in mice treated with EcNc^Δ*lon*/Δ*ompT*/LLO+^ nAg^19^ in comparison to treatment with EcNc^Δ*lon*/Δ*ompT*^ or PBS (**Fig. 3d**).

Beyond induction of tumor antigen–specific T cell responses, treatment with EcNc^Δ*lon*/Δ*ompT*/LLO+^ nAg^19^ resulted in reduced frequencies of tumor-resident immunosuppressive PD-L1^+^Ly6G^+^ polymorphonuclear cells (PMNs) and PD-L1^+^F4/80^+^ macrophages (**Fig. 3e**)^54,55^. Bacteria-treated groups further displayed reduced numbers and frequencies of Foxp3^+^CD4^+^ regulatory T cells and MHC-II^lo^F4/80^+^ tumor-associated macrophages (**Fig. 3f, Extended Data Fig. 7d–e**), two cell populations known for their roles in inhibiting anti-tumor immunity^56,57^. Moreover, myeloid immunophenotyping revealed a reduction of PD-L1 on cDC1 and cDC2 populations within TDLNs of the neoantigen therapeutic treated group (**Extended Data Fig. 7f**), which has been shown to facilitate and sustain anti-tumor immunity^58,59^. In summary, intravenously delivered microbial neoantigen therapeutics sustain neoantigen production and availability in lymphoid tissue *in vivo*, stimulate both neoantigen-specific and broad adaptive immunity, and reduce immunosuppression within the TME, shaping a more effective environment for productive anti-tumor immunity (**Fig. 3g**).

### Microbial anti-tumor vaccination in melanoma

Neoantigens are generally unique to the individual patient^60^, thus vaccination platforms must be able to flexibly incorporate and deliver diverse sets of neoantigens based on the unique mutations present in a particular tumor. To evaluate the suitability of our engineered microbial platform in this regard, we performed paired exome and transcriptome sequencing on a second, more aggressive tumor cell type (B16F10 melanoma) grown orthotopically in C57BL/6 mice and designed tumor-specific therapeutics (**Fig. 4a**). We applied an equivalent neoantigen prediction algorithm as performed for CT26 and identified numerous putative B16F10-specific neoantigens, including many that had previously been validated by others^23,61^ (**Extended Data Fig. 8a**). A set of 7 constructs were devised from neoantigens of varying imputed MHC-I and MHC-II affinities, with each construct containing 6 unique predicted neoantigens (**Extended Data Table 3**) and confirmed to be robustly expressed by EcNc^Δ*lon*/Δ*ompT*/LLO+^ (**Fig. 4a**, **Extended Data Fig. 8b**).

**Figure 4.**
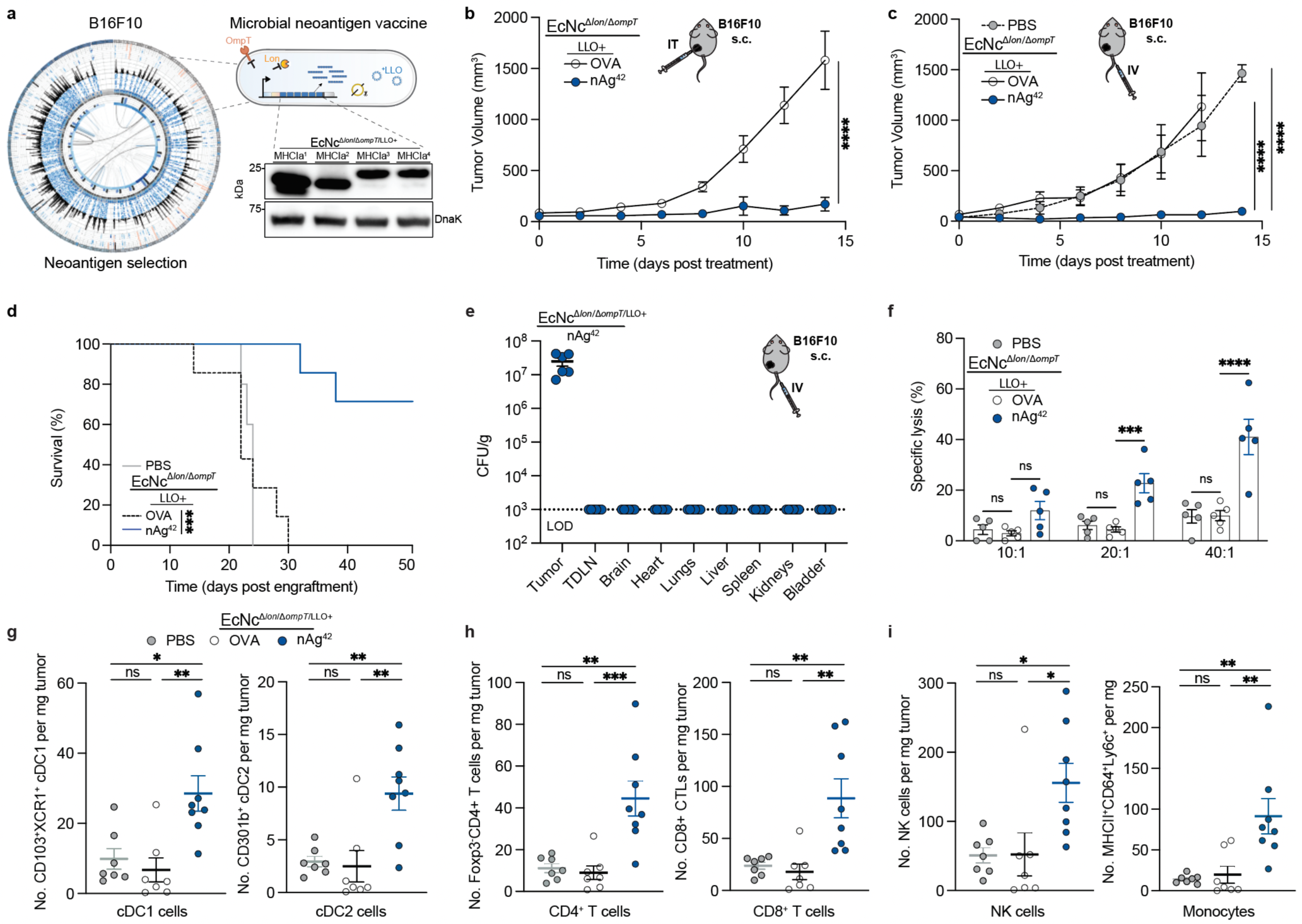
Efficacy of microbial anti-tumor vaccines in orthotopic melanoma. **a**, Design of microbial neoantigen therapeutics for melanoma. Left: Circos plot depicting mutanome of B16F10. Right: immunoblot of B16F10 neoantigen constructs. **b**–**e**, **g**–**i**, C57BL/6 mice with established hind-flank B16F10 melanoma tumors were treated starting 9 days after tumor engraftment. **b**, Mice received an intratumoral injection of EcNc^Δ*lon*/Δ*ompT*/LLO+^ OVA or EcNc^Δ*lon*/Δ*ompT*/LLO+^ nAg^42^ every 3–5 days. Tumor growth curves (*n* = 7 mice per group, \*\*\*\**P* < 0.0001, ns = not significant (*P* > 0.05), two-way ANOVA with Šídák’s multiple comparisons test). **c**, Mice received an intravenous injection of either PBS, EcNc^Δ*lon*/Δ*ompT*/LLO+^ OVA or EcNc^Δ*lon*/Δ*ompT*/LLO+^ nAg^42^ every 3–5 days. Tumor growth curves (*n* = 5–7 mice per group, \*\*\*\**P* < 0.0001, ns = not significant (*P* > 0.05), two-way ANOVA with Tukey’s multiple comparisons test). **d**, Kaplan-Meier survival curves for orthotopic B16F10 tumor-bearing mice from (**c**) (*n* = 5–7 mice per group, \*\*\**P* = 0.0001, Log-rank Mantel-Cox test). **e**, Mice (*n* = 6 per group) received intravenous injection of the 7-strain combination EcNc^Δ*lon*/Δ*ompT*/LLO+^ nAg^42^. Microbial tissue burden quantified by CFU per gram of tissue (CFU/g), LOD = 1 × 10^3^ CFU/g. **f**, Naïve, tumor free C57BL/6 mice were vaccinated intravenously with the designated treatment. Specific lysis of B16F10-Luc cells by purified splenic T cells from mice (*n* = 5 per group) at specified effector-to-target cell (E:T) ratios (\*\*\**P* = 0.001, \*\*\*\**P* < 0.0001, ns = not significant (*P* > 0.05), one-way ANOVA with Tukey’s multiple comparisons test). **g**–**i**, Mice received intravenous injection of either PBS, EcNc^Δ*lon*/Δ*ompT*/LLO+^ OVA or EcNc^Δ*lon*/Δ*ompT*/LLO+^ nAg^42^ on day 9 and 12 post orthotopic B16F10 engraftment. **g**, Left: Number of CD103^+^XCR1^+^ cDC1 cells per mg tumor (*n* = 7-8 mice per group, \**P* = 0.0103, \*\**P* = 0.0030, ns = not significant (*P* > 0.05), one-way ANOVA with Tukey’s multiple comparisons test). Right: Number of CD310b^+^ cDC2 cells per mg tumor (*n* = 7–8 mice per group, \*\**P* = 0.0038, \*\**P* = 0.0064, ns = not significant (*P* > 0.05), one-way ANOVA with Tukey’s multiple comparisons test). **h**, Left: Number of Foxp3^-^CD4^+^ T cells per mg tumor (*n* = 7-8 mice per group, \*\**P* = 0.0015, \*\*\**P* = 0.0008, ns = not significant (*P* > 0.05), one-way ANOVA with Tukey’s multiple comparisons test). Right: Number of CD8^+^ cytotoxic T cells per mg tumor (*n* = 7–8 mice per group, \*\**P* = 0.0022, \*\**P* = 0.0047, ns = not significant (*P* > 0.05), one-way ANOVA with Tukey’s multiple comparisons test). **i**, Left: Number of NK1.1^+^ NK cells per mg tumor (*n* = 7-8 mice per group, \**P* = 0.0243, \**P* = 0.0224, ns = not significant (*P* > 0.05), one-way ANOVA with Tukey’s multiple comparisons test). Right: Number of MHCII^+^CD64^+^Ly6c^+^ monocytes per mg tumor (*n* = 7–8 mice per group, \*\**P* = 0.0041, \*\**P* = 0.0073, ns = not significant (*P* > 0.05), one-way ANOVA with Tukey’s multiple comparisons test). **b**–**e**, **d**–**i**, Data are mean ± s.e.m.

We then sought to test the anti-tumor efficacy of our therapeutics against advanced B16F10 tumors. When established orthotopic tumors were injected with microbial therapeutics intratumorally, tumors grew progressively after treatment with EcNc^Δ*lon*/Δ*ompT*/LLO+^ expressing the strong irrelevant xenoantigen ovalbumin (OVA), whereas treatment with the equal-parts combination of all 7 construct expressing strains – encompassing 42 unique B16F10 neoantigens (nAg^42^) – significantly repressed growth in EcNc^Δ*lon*/Δ*ompT*/LLO+^ nAg^42^ treated tumors over the same time course (**Fig. 4b**). Similarly, intravenous treatment with EcNc^Δ*lon*/Δ*ompT*/LLO+^ nAg^42^ potently restrained orthotopic tumor growth, with 72% of nAg^42^ treated mice alive 50 days post-tumor engraftment, while all control group mice succumbed to malignancy by day 24 or 30 (**Fig. 4c, d, Extended Data Fig. 8c**). Treatment with intravenous microbial vaccines again induced no significant body weight change as compared to PBS treated mice (**Extended Data Fig. 8d**).

To evaluate the tissue biodistribution of the microbial neoantigen vaccines after systemic administration in this setting, we surveyed organs after intravenous injection of EcNc^Δ*lon*/Δ*ompT*/LLO+^ nAg^42^. As we observed for BALB/c mice with CT26 tumors, live microbial vectors specifically colonized the B16F10 tumor at high density without detectable presence in any other organs examined (**Fig. 4e**). To confirm that the microbial B16F10 neoantigen vaccine generated T cells capable of direct tumor cell killing, we treated tumor-free C57BL/6 mice intravenously with microbial therapeutics and co-incubated purified splenic T cells with B16F10 tumor cells *in vitro*. Indeed, T cells from mice treated intravenously with EcNc^Δ*lon*/Δ*ompT*/LLO+^ nAg^42^ but not EcNc^Δ*lon*/Δ*ompT*/LLO+^ OVA demonstrated enhanced killing of B16F10 tumor cells (**Fig. 4f**). These data verify tumor-specific colonization and antigen-specific T cell induction by microbial neoantigen vaccines in B16F10 melanoma.

### Microbial vectors modulate anti-tumor immunity and metastases in melanoma

To characterize the immunologic changes associated with anti-tumor efficacy in this alternate model, we performed immunophenotyping of the orthotopic B16F10 tumors 8 days post intravenous microbial treatment. Tumors treated intravenously with EcNc^Δ*lon*/Δ*ompT*/LLO+^ nAg^42^ as compared to controls had significantly higher numbers and frequencies of cDC1 and cDC2 cells, conventional CD4^+^ and cytotoxic CD8^+^ infiltrating T cells, NK cells, and inflammatory monocytes (**Fig. 4g**–**i****, Extended Data Fig. 8e–g**).

Analyses of the intratumoral lymphoid compartment revealed enhanced expression of CD69 on Foxp3^-^CD4^+^ and CD8^+^ TILs, and significantly increased frequencies of IFN-γ secreting conventional Foxp3^-^CD4^+^ and cytotoxic CD8^+^ TILs after restimulation with PMA and ionomycin in EcNc^Δ*lon*/Δ*ompT*/LLO+^ nAg^42^–treated tumors, indicating enhanced T cell activation and effector cytokine production within the TME (**Fig. 5a**, **Extended Data Fig. 9a**). Tumor-infiltrating Foxp3^−^ CD4^+^ and CD8^+^ T cells and NK cells also expressed significantly higher levels of granzyme-B after EcNc^Δ*lon*/Δ*ompT*/LLO+^ nAg^42^ treatment, suggestive of amplified cytolytic function (**Fig. 5b**). Consistent with enduring activity of anti-tumor immunity, we also observed higher levels of proliferating tumor-infiltrating CD4^+^ and CD8^+^ T cells, and NK cells as assessed by Ki-67 staining (**Extended Data Fig. 9b**).

**Figure 5.**
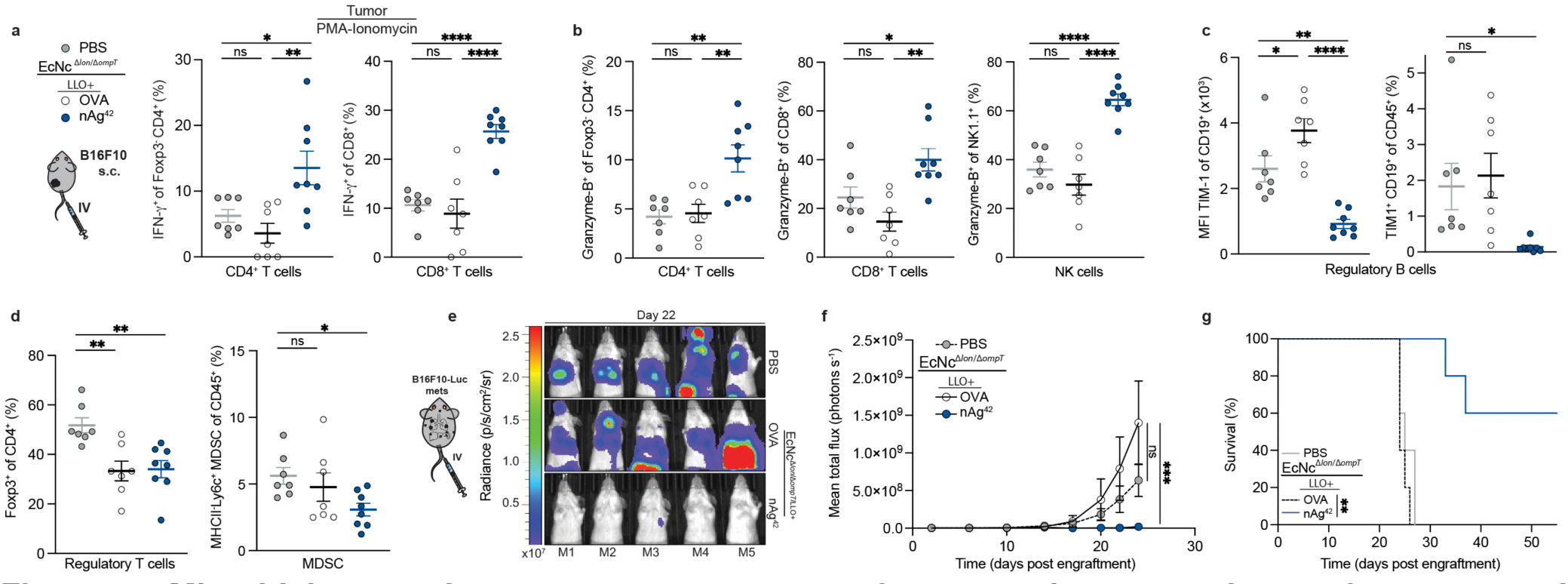
Microbial neoantigen vectors restructure the tumor immune microenvironment and suppress established metastatic melanoma. **a–d**, Mice received an intravenous injection of either PBS, EcNc^Δ*lon*/Δ*ompT*/LLO+^ OVA or EcNc^Δ*lon*/Δ*ompT*/LLO+^ nAg^42^ on day 9 and 12 post orthotopic B16F10 engraftment. **a**, Tumor-infiltrating lymphocytes were stimulated with PMA and ionomycin in the presence of brefeldin A. Left: experimental schematic. Middle: Frequency of IFN-γ_+_ Foxp3^-^CD4^+^ post-stimulation (*n* = 7–8 mice per group, \**P* = 0.0335, \*\**P* = 0.0040, ns = not significant (*P* > 0.05), one-way ANOVA with Tukey’s multiple comparisons test). Right: frequency of IFN-γ_+_ CD8^+^ T cells in tumors (*n* = 7–8 mice per group, \*\*\*\**P* < 0.0001, ns = not significant (*P* > 0.05), one-way ANOVA with Tukey’s multiple comparisons test). **b**, Left: Frequency of Granzyme-B^+^ Foxp3^-^CD4^+^ in tumors (*n* = 7–8 mice per group, \*\**P* = 0.0024, \*\**P* = 0.0041, ns = not significant (*P* > 0.05), one-way ANOVA with Tukey’s multiple comparisons test). Middle: Frequency of Granzyme-B^+^ CD8^+^ T cells in tumors (*n* = 7–8 mice per group, \**P* = 0.0495, \*\**P* = 0.0014, ns = not significant (*P* > 0.05), one-way ANOVA with Tukey’s multiple comparisons test). Right: Frequency of Granzyme-B^+^ NK1.1^+^ NK cells in tumors (*n* = 7–8 mice per group, \*\*\*\**P* < 0.0001, ns = not significant (*P* > 0.05), one-way ANOVA with Tukey’s multiple comparisons test). **c**, Left: Median fluorescence intensity (MFI) of TIM-1 on CD19^+^ B cells in tumors (*n* = 7–8 mice per group, \**P* = 0.0457, \*\**P* = 0.0029, \*\*\*\**P* < 0.0001, one-way ANOVA with Tukey’s multiple comparisons test). Right: Frequency of TIM-1^+^CD19^+^ B cells of CD45^+^ cells in tumors (*n* = 7-8 mice per group, \**P* = 0.0442, ns = not significant (*P* > 0.05), one-way ANOVA with Dunnett’s multiple comparisons test). **d**, Left: Frequency of Foxp3^+^CD4^+^ T cells of CD4^+^ cells in tumors (*n* = 7-8 mice per group, \*\**P* = 0.0035, \*\**P* = 0.0038, one-way ANOVA with Dunnett’s multiple comparisons test). Right: Frequency of MHCII^-^Ly6c^+^ MDSC of CD45^+^ cells in tumors (*n* = 7-8 mice per group, \**P* = 0.0440, ns = not significant (*P* > 0.05), one-way ANOVA with Dunnett’s multiple comparisons test). **e**–**g**, Mice received intravenous injection of either PBS, EcNc^Δ*lon*/Δ*ompT*/LLO+^ OVA or EcNc^Δ*lon*/Δ*ompT*/LLO+^ nAg^42^ every 3–5 days starting 2 days after intravenous injection of B16F10-Luc cells. **e**, Representative images of lung metastases luminescence in each mouse (M1–M5) per group on day 22 post engraftment. **f**, Mean total flux from systemic metastases (*n* = 5 mice per group, \*\*\**P* > 0.0003, ns = not significant (*P* > 0.05), two-way ANOVA with Dunnett’s multiple comparisons test). **g**, Kaplan-Meier survival curve for mice with B16F10-Luc systemic metastases (*n* = 5 mice per group, \*\**P* = 0.0015, Log-Rank Mantel-Cox test). **a**– **d, f, g**, Data are mean ± s.e.m.

In addition to the enhanced activation of tumor-infiltrating T and NK cells, treatment with EcNc^Δ*lon*/Δ*ompT*/LLO+^ nAg^42^ significantly reduced TIM-1 expression by tumor-infiltrating CD19^+^ B cells and thus the frequency of regulatory TIM-1^+^ B cells – an important immunosuppressive cell population in the B16F10 model^62^ – and increased B cell proliferation (**Fig. 5c**, **Extended Data Fig. 9c**). Moreover, EcNc^Δ*lon*/Δ*ompT*/LLO+^ nAg^42^ vaccination reduced the frequency of immunosuppressive Foxp3^+^ regulatory T cells, myeloid-derived suppressor cells (MDSC), and MHC-II^lo^ macrophages within tumors (**Fig. 5d**, **Extended Data Fig. 9d**). Infiltrating monocytes and dendritic cells in EcNc^Δ*lon*/Δ*ompT*/LLO+^ nAg^42^–treated tumors exhibited increased expression of MHC-II (**Extended Data Fig. 9e**), suggestive of enhanced antigen presentation capacity of these cell types within tumors. Overall, these data demonstrate that intravenous microbial tumor neoantigen vaccination mediates immunologic restructuring within the melanoma TME, recruiting APCs and activating NK cells, and CD4^+^ and CD8^+^ T cells while diminishing central immunosuppressive cell populations.

Given the robust anti-tumor efficacy induced by vaccination in orthotopic B16F10, we investigated the efficacy of EcNc^Δ*lon*/Δ*ompT*/LLO+^ nAg^42^ in established, systemic B16F10-Luc metastases. Whereas systemic metastases rapidly progressed in PBS or EcNc^Δ*lon*/Δ*ompT*/LLO+^ OVA treated mice, EcNc^Δ*lon*/Δ*ompT*/LLO+^ nAg^42^ strongly inhibited metastatic growth (**Fig 5. e–f, Extended Data Fig. 10a, b**). Treatment with EcNc^Δ*lon*/Δ*ompT*/LLO+^ nAg^42^ significantly extended survival, with 60% of mice surviving to 55 days with no detectable metastases, whereas all control treated mice had died by day 27 (**Fig 5. g**). Again, treatment was well-tolerated, with no significant weight change relative to PBS (**Extended Data Fig. 10c**). These data demonstrate that the microbial tumor neoantigen vaccination system stimulates robust and effective anti-tumor immunity *in vivo* after intravenous administration in established, systemic metastatic melanoma.

## Discussion

Through microbial engineering, we couple the tumor-homing and immunostimulatory nature of bacteria with precise instructions for coordinated adaptive immunity toward tumor neoantigens. Optimization of neoantigen constructs, genetic editing and cryptic plasmid removal, and expression of LLO for phagocytic escape together achieve an anti-tumor vaccination platform capable of delivering diverse arrays of neoantigens to mediate control and eradication of advanced primary and metastatic solid tumors.

We show that across distinct tumor models and genetic backgrounds, the anti-tumor effect of vaccination is accompanied by broad modulation of the immune compartment within the TME. The coordinated regulation of APCs, reduction of immunosuppressive myeloid, regulatory T and B cell populations, and activation of NK, and CD4^+^ and CD8^+^ T effector cells indicates the advantage of precisely engineered microbial platforms as next generation anti-tumor vaccines which align multiple arms of immunity^21^.

Tumor- and patient-specific modifications to this microbial platform can be conducted rapidly due to ease of design and delivery of unique neoantigen cassettes. We found that composite sets encompassing both predicted MHC-I and MHC-II binding neoantigens were able to mediate anti-tumor efficacy. This is in agreement with the critical role of both CD4^+^ and CD8^+^ T cells in effective anti-tumor immunity, and the expanding set of verified MHC-I and MHC-II dependent neoantigens recognized across murine and human tumors alike^19,23,24,61,63,64^. However, further investigation is necessary to elucidate the optimal characteristics of incorporated neoantigen sets and the neoantigen-intrinsic factors underlying efficacy in the system created here. Additional programming of tumor neoantigen constructs, and rational incorporation of other immunotherapeutics through synthetic modification, may synergistically achieve reliable eradication of advanced solid tumors and associated metastases through precision cancer immunotherapy using living anti-tumor vaccines.

## Extended Data

**Extended Data Figure 1:**
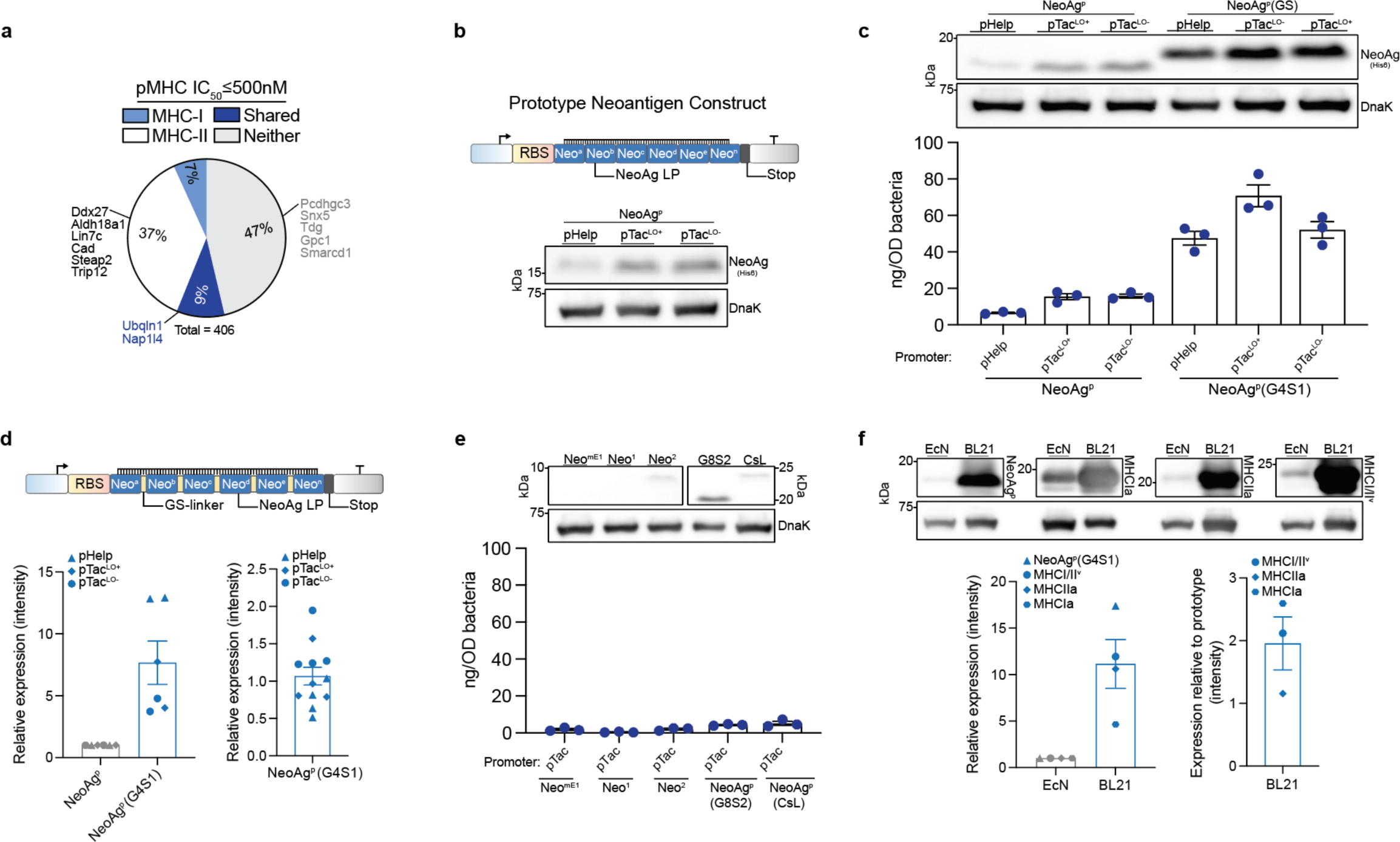
Neoantigen prediction and synthetic construct design. **a**, Percentage of predicted CT26 neoantigens containing mutant-epitope(s) with ≤500nM MHC-I affinity (MHC-I), MHC-II affinity (MHC-II), both MHC-I and MHC-II affinity (Shared), or no epitope meeting affinity criteria (Neither). Previously validated neoantigens within the set are labeled. **b**, Upper: prototype neoantigen construct design, Lower: immunoblot of EcN expressing prototype neoantigen constructs. **c**, Upper: immunoblot of EcN expressing prototype neoantigen constructs with or without GS-linkers. Lower: ELISA quantification of neoantigen construct in soluble fraction with or without GS-linkers in DH5α (*n* = 3 per group). NeoAg^p^ = prototype neoantigen construct, G4S1 = 5-mer GS-linker, pTac^LO-^ = pTac without Lac operator; pTac^LO+^ = with Lac operator. **d**, Upper: neoantigen construct design with GS-linkers, Lower-left: Relative immunoblot chemiluminescent intensity for prototype construct with or without interspersing glycine-serine linkers (*n* = 6 per group). Lower-right: relative expression of prototype neoantigen construct with GS-linkers under selected promoters (*n* = 12 samples). **e**, Upper: immunoblot of EcN expressing alternate prototype neoantigen constructs. Lower: ELISA quantification of alternative neoantigen construct in soluble fraction (*n* = 3 per group). Neo^mE1^ = minimal epitope, Neo^1^ = 1 neoantigen LP in construct, Neo^2^ = 2 neoantigen LP in construct, G8S2 = 10-mer GS-linker, CsL = immunoprotease sensitive linker. **f**, Upper: Immunoblot of neoantigen constructs (NeoAg^p^, MHCIa, MHCIIa, MHCI/II^v^), expressed in BL21 or EcN. Lower-left: relative immunoblot chemiluminescent intensity for neoantigen construct expression in EcN vs. BL21 (*n* = 4 per group), Lower-right: relative immunoblot chemiluminescent intensity of predicted neoantigen constructs vs. prototype in BL21 (*n* = 3 samples). **c**–**f**, Data are mean ± s.e.m.

**Extended Data Figure 2:**
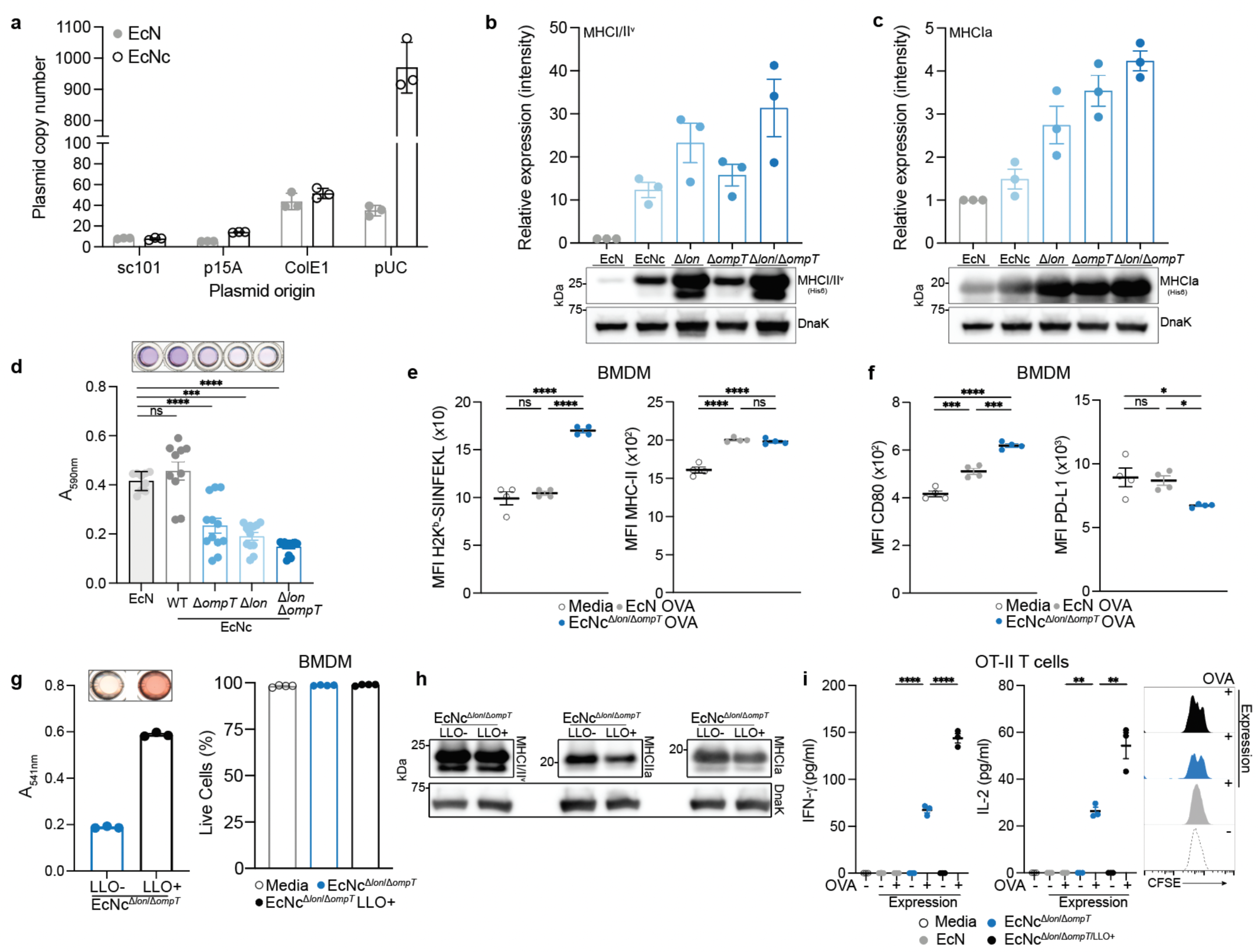
Evaluation of microbial tumor neoantigen vaccine functioning and immunologic activity *in vitro*. **a**, Plasmid copy number in wildtype EcN or cryptic plasmid cured EcNc (*n* = 3 per group). **b**, Upper: relative immunoblot chemiluminescent intensity of synthetic neoantigen construct MHCI/II^v^ expression in wildtype EcN vs. derivative strains (*n* = 3 replicates), Lower: representative immunoblot of construct MHCI/II^v^ expressed in wildtype EcN and derivative strains. **c**, Upper: relative immunoblot chemiluminescent intensity of synthetic neoantigen construct MHCIa expression in wildtype EcN vs. derivative strains (*n* = 3 replicates), Lower: representative immunoblot of construct MHCIa expressed in wildtype EcN and derivative strains. **d,** Biofilm formation quantified for wildtype EcN, and derivative strains by crystal violet stain assay. (\*\*\*\**P*<0.0001, ns = not significant *P* > 0.05, One-way ANOVA with Tukey’s multiple comparison test, *n* = 9-12 per group). **e**, Median fluorescence intensity (MFI) of H2k^b^-SIINFEKL complex and MHCII, or **f**, CD80 and PD-L1 for BMDM incubated with the indicated live microbial strain or culture media for 6 hours (\**P* = 0.0231, \**P* = 0.0414, \*\*\**P* = 0.0002, \*\*\**P* = 0.0005, \*\*\*\**P* < 0.0001, one-way ANOVA with Tukey’s multiple comparisons test, *n* = 4 per group). **g**, Left: sheep red blood cells (RBCs) were incubated with lysate from EcNc^Δ*lon*/Δ*ompT*^ with (LLO+) or without (LLO-) cytosolic LLO expression. Absorbance at 541nm (*n* = 3 per group). Right: percentage of live BMDM after incubation with indicated live microbial strain or control for 6 hours (*n* = 4 per group). **h**, Immunoblot depicting expression of neoantigen constructs MHCIa, MHCIIa, and MHCI/II^v^ in EcNc^Δ*lon*/Δ*ompT*^ with (LLO+) or without (LLO-) co-expression of cytosolic LLO. **i**, Naïve OT-II T cells were incubated with BMDC’s pulsed with the indicated condition. Left: IFN-γ quantification in supernatant of OT-II cultures (\*\*\*\**P* < 0.0001, one-way ANOVA with Tukey’s multiple comparisons test, *n* = 3 replicates per group), Middle: IL-2 quantification in supernatant of OT-II culture (\*\*\*\**P* < 0.0001, one-way ANOVA with Tukey’s multiple comparisons test, *n* = 3 replicates per group). Right: representative histogram depicting CFSE dilution of stimulated OT-II T cells. **a**–**i**, Data are mean ± s.e.m.

**Extended Data Figure 3.**
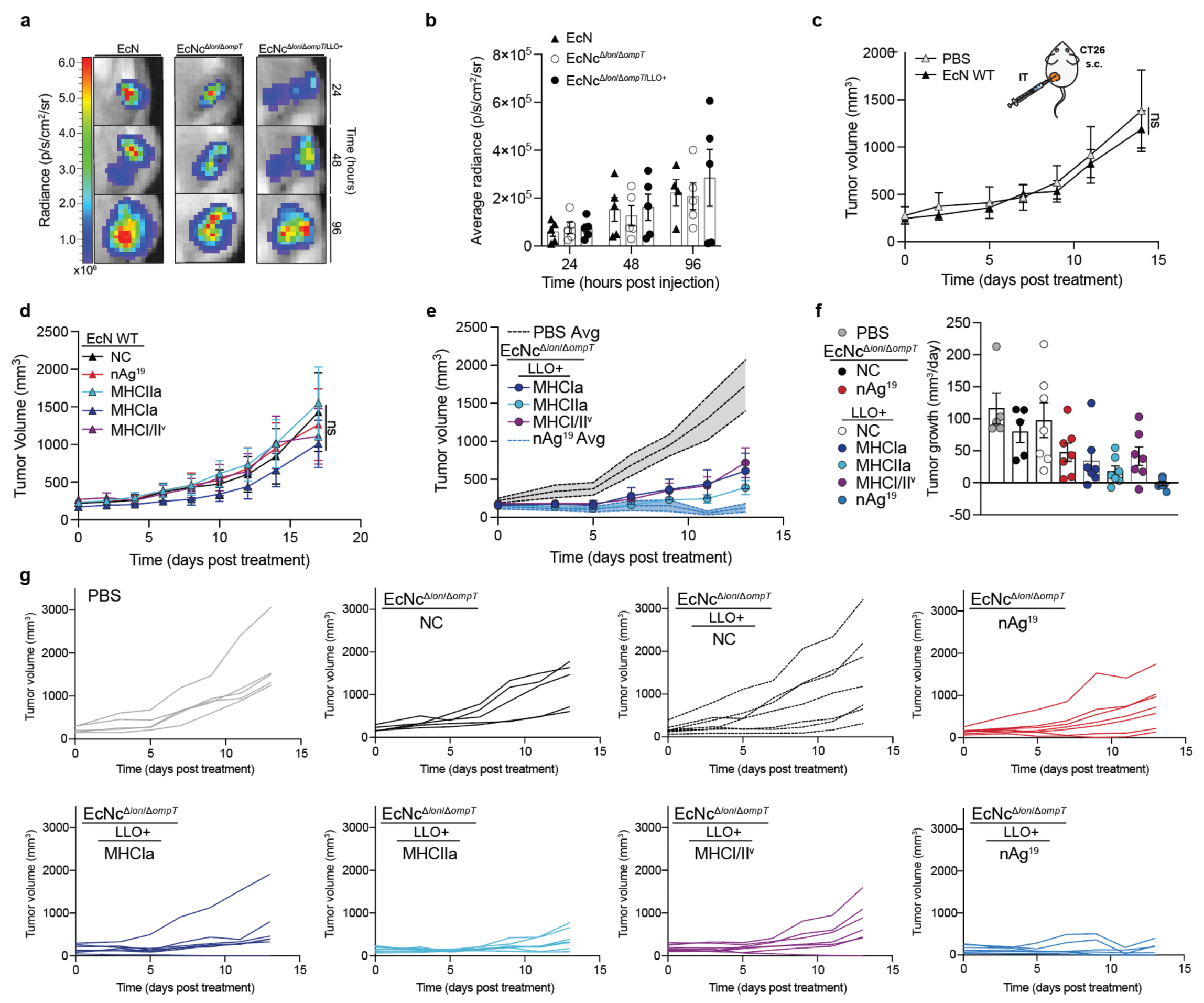
Characterization of intratumoral treatment with microbial tumor neoantigen vaccines. **a**–**g,** BALB/c mice with established hind-flank CT26 tumors were treated when average tumor volumes were ∼150-200mm^3^. **a**-**b**, Mice received a single intratumoral injection of EcN WT, EcNc^Δ*lon*/Δ*ompT*^, or EcNc^Δ*lon*/Δ*ompT*/LLO+^. **a**, Representative image of tumors colonized by microbes with a genome-integrated luminescence cassette. **b**, Average radiance of microbe colonized tumors in designated groups post intratumoral injection (*n* = 4-5 tumors per group). **c,** Mice received intratumoral injections of PBS or EcN WT. Tumor growth curves (*n* = 5 mice per group, ns = not significant (*P* > 0.05), two-way ANOVA with Tukey’s multiple comparisons test). **d**, Mice received intratumoral injections of EcN WT without therapeutic expression (NC), expressing either construct MHCIa, MHCIIa, or MHCI/II^v^, or EcN nAg^19^. Tumor growth curves (*n* = 5 mice per group, ns = not significant (*P* > 0.05), two-way ANOVA with Tukey’s multiple comparisons test). **e** and **f**, Mice (*n =* 5-7 per group) received a single intratumoral injection of PBS, EcNc^Δ*lon*/Δ*ompT*^ NC or EcNc^Δ*lon*/Δ*ompT*^ nAg^19^, or EcNc^Δ*lon*/Δ*ompT*/LLO+^ expressing either construct MHCIa, MHCIIa, or MHCI/II^v^, or EcNc^Δ*lon*/Δ*ompT*/LLO+^ nAg^19^. **e**, Tumor growth curves, and **f**, Tumor growth rate (mm^3^ per day) for the respective treatment group. **g**, Individual tumor trajectories (*n* = 5-7 mice per group) after intratumoral treatment with PBS or indicated microbial therapeutic. **b**–**f**, Data are mean ± s.e.m.

**Extended Data Figure 4.**
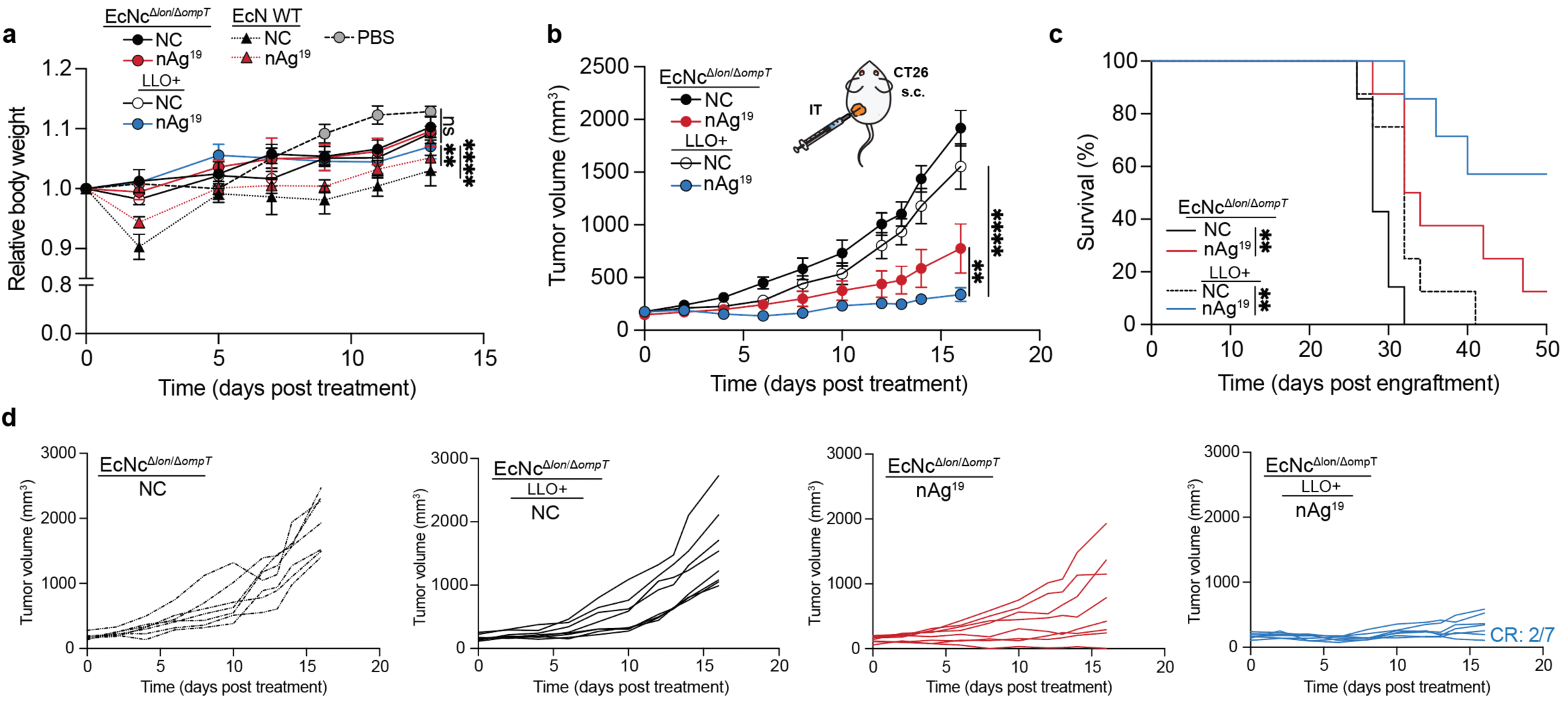
Comparative profile of intratumoral treatment with engineered microbial neoantigen vaccines. **a**–**d,** BALB/c mice with established hind-flank CT26 tumors were treated when average tumor volume was 150-200mm^3^. **a**, Mice received intratumoral injection of wildtype EcN, EcNc^Δ*lon*/Δ*ompT*^ or EcNc^Δ*lon*/Δ*ompT*/LLO+^ strain combination expressing the 3 neoantigen constructs (nAg^19^), or without neoantigen expression (NC) on day 0. Relative body weight of CT26 tumor-bearing mice (*n* = 5-7 mice per group, \*\**P* = 0.0034, \*\*\*\**P* <0.0001, ns = not significant (*P* > 0.05), two-way ANOVA with Tukey’s multiple comparisons test). **b**, Mice received intratumoral injection on day 0 and 8. Tumor growth curves (*n* = 7-8 mice per group, \*\**P* = 0.0020, \*\*\*\**P* <0.0001, two-way ANOVA with Tukey’s multiple comparisons test). **c**, Kaplan-Meier survival curves for CT26 tumor-bearing mice (*n* = 7-8 mice per group, \*\**P* = 0.0061, \*\**P* = 0.0076, Log-rank Mantel-Cox test). **d**, Individual tumor trajectories (*n* = 7–8 mice per group) after intratumoral treatment with indicated microbial strain. **a**–**c,** Data are mean ± s.e.m.

**Extended Data Figure 5.**
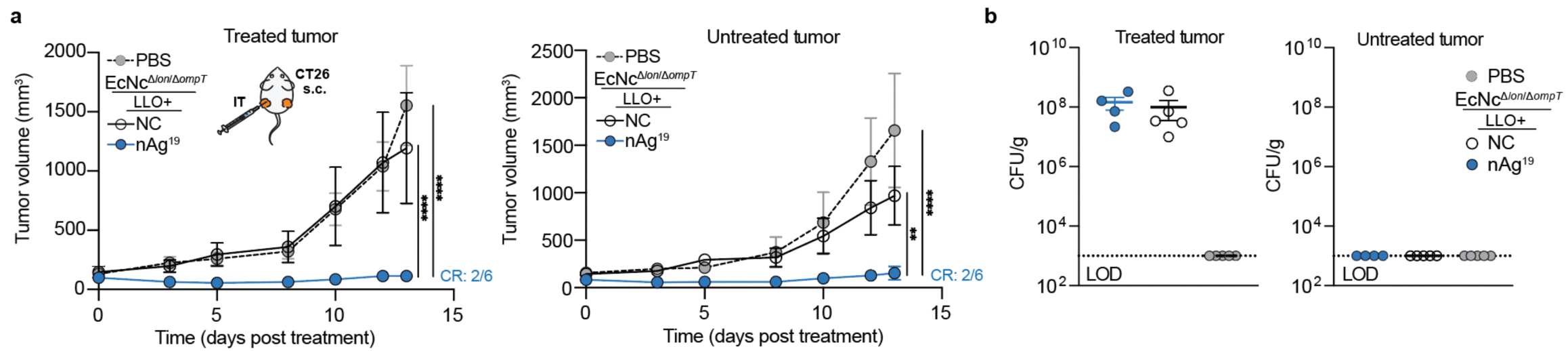
Induction of systemic anti-tumor immunity after treatment of a single tumor. **a** and **b**, BALB/c mice were implanted with CT26 cells on both hind flanks. When average tumor volumes were ∼100-150mm^3^ mice received an intratumoral injection of PBS, EcNc^Δ*lon*/Δ*ompT*/LLO+^ (NC), or EcNc^Δ*lon*/Δ*ompT*/LLO+^ nAg^19^ into a single tumor. **a**, Tumor growth curves (*n* = 5-6 mice per group, \*\**P* = 0.0014, \*\*\*\**P* < 0.0001, two-way ANOVA with Tukey’s multiple comparisons test). **b**, CFU/g of tumor (*n* = 5-6 mice per group), LOD 1 × 10^3^ CFU. **a** and **b,** Data are mean ± s.e.m.

**Extended Data Figure 6.**
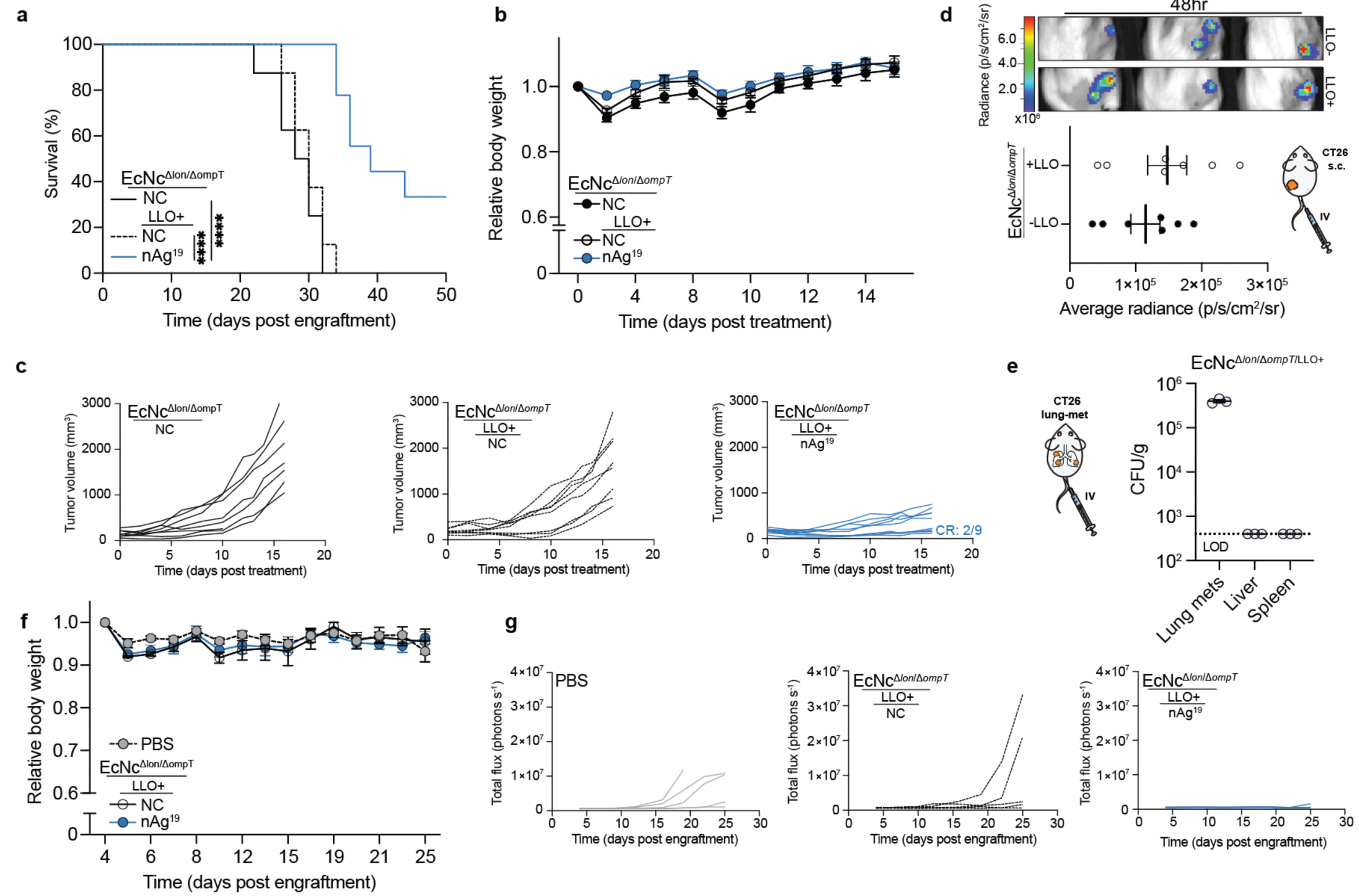
Intravenous engineered microbial treatment in primary and metastatic solid tumors. **a**–**c**, BALB/c mice with established hind-flank CT26 tumors were treated when average tumor volume was ∼150-200mm^3^. **a**, Kaplan-Meier survival curves for CT26-tumor-bearing mice treated with indicated therapeutic (*n* = 8-9 mice per group, \*\*\*\**P* < 0.0001, \*\**P* = 0.0021, Log-rank Mantel-Cox test). **b**, Relative body weight of mice (*n* = 8-9 per group) after intravenous treatment with indicated microbial therapeutic on day 0 and 8. **c**, Individual tumor trajectories (*n* = 8-9 mice per group) after intravenous treatment with the indicated microbial therapeutic. **d**, Mice received an intravenous injection of EcNc^Δ*lon*/Δ*ompT*^ or EcNc^Δ*lon*/Δ*ompT*/LLO+^. Upper: representative luminescent signature of tumors colonized with EcNc^Δ*lon*/Δ*ompT*^(-LLO) or EcNc^Δ*lon*/Δ*ompT*/LLO+^ (+LLO), 48 hours post-injection. Lower: average radiance of colonized tumors (*n* = 7 mice per group). **e**–**g**, BALB/c mice were injected intravenously with CT26-Luc cells. Every 3-5 days, mice (*n* = 5 per group) received intravenous injection of either PBS, EcNc^Δ*lon*/Δ*ompT*/LLO+^ without therapeutic (NC), or the 3 neoantigen-construct expressing strain combination nAg^19^. **e**, Microbial tissue burden quantified as CFU/g tissue, LOD 4×10^2^ CFU (*n* = 3 mice). **f**, Relative body weight of mice (*n* = 5 per group) after intravenous treatment with indicated therapeutic. **g**, Individual lung metastases luminescence trajectories (*n* = 5 mice per group). **a**–**g**, Data are mean ± s.e.m.

**Extended Data Figure 7.**
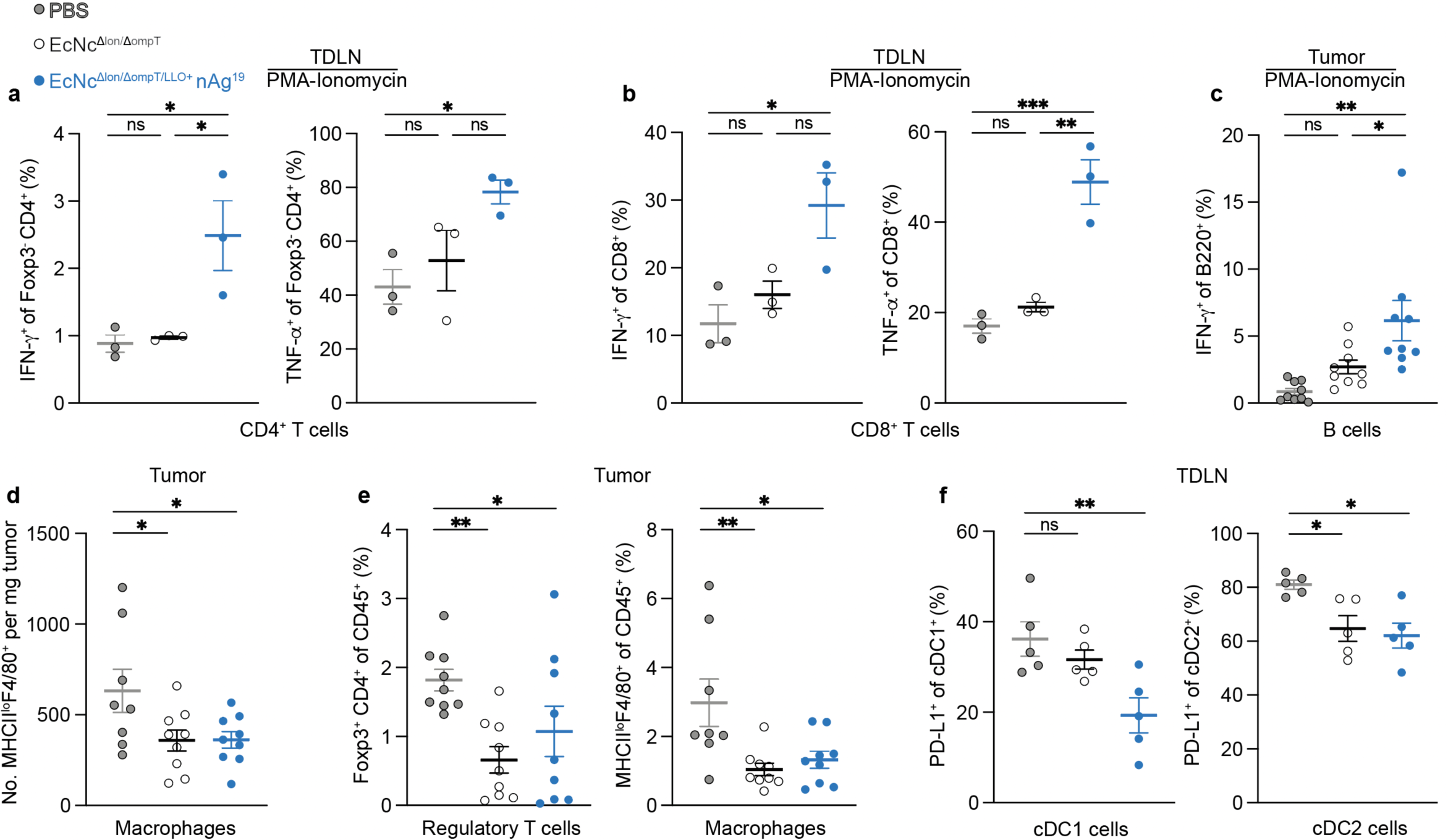
Modulation of the tumor-immune microenvironment by engineered microbial tumor neoantigen vaccines. **a**–**c**, BALB/c mice with established hind-flank CT26 tumors received an intravenous injection of indicated therapeutic or control. 8 days after treatment, tumors and TDLN’s were extracted. **a**–**b**, Lymphocytes from TDLNs were stimulated *ex vivo* with PMA and ionomycin in the presence of brefeldin A. **a**, Left: Frequency of IFN-γ_+_ Foxp3^-^CD4^+^ post-stimulation (*n* = 3 mice per group, \**P* = 0.0313, \**P* = 0.0246, ns = not significant (*P* > 0.05), One-way ANOVA with Tukey’s multiple comparisons test). Right: Frequency of TNF-α_+_ Foxp3^-^CD4^+^ T cells post-stimulation (*n* = 3 mice per group, \**P* = 0.0445, ns = not significant (*P* > 0.05), One-way ANOVA with Tukey’s multiple comparisons test). **b**, Left: Frequency of IFN-γ_+_ CD8^+^ post-stimulation (*n* = 3 mice per group, \**P* = 0.0257, ns = not significant (*P* > 0.05), One-way ANOVA with Tukey’s multiple comparisons test). Right: Frequency of TNF-α_+_ CD8^+^ T cells post-stimulation (*n* = 3 mice per group, \*\**P* = 0.0017, \*\*\**P* = 0.0008, ns = not significant (*P* > 0.05), One-way ANOVA with Tukey’s multiple comparisons test). **c**, Frequency of IFN-γ_+_ B220^+^ B cells post-stimulation (*n* = 9 mice per group, \**P* = 0.0351, \*\**P* = 0.0010, ns = not significant (*P* > 0.05), One-way ANOVA with Tukey’s multiple comparisons test). **d**, Number of MHC-II^lo+^F4/80^+^ CD11b^+^ macrophages per mg tumor (*n* = 8-9 mice per group, \**P* = 0.0385, \**P* = 0.0407, one-way ANOVA with Dunnett’s multiple comparisons test). **e**, Left: Frequency of FoxP3^+^CD4^+^ regulatory T cells in tumors (*n* = 9 mice per group, \**P* = 0.0491, \*\**P* = 0.0072, one-way ANOVA with Holm-Šídák’s multiple comparisons test), Right: Frequency of MHCII^lo^F4/80^+^CD11b^+^ macrophages in tumors (*n* = 8-9 mice per group, \**P* = 0.0173, \*\**P* = 0.0057, one-way ANOVA with Dunnett’s multiple comparisons test). **f**, Left: Percentage PD-L1^+^ of cDC1 in TDLN (*n* = 5 mice per group, \*\**P* = 0.0074, ns = not significant (*P* > 0.05), one-way ANOVA with Dunnett’s multiple comparisons test), Right: Percentage PD-L1^+^ of cDC2 in TDLN (*n* = 5 mice per group, \**P* = 0.0103, \**P* = 0.0244, one-way ANOVA with Dunnett’s multiple comparisons test). **a**–**f**, Data are mean ± s.e.m.

**Extended Data Figure 8.**
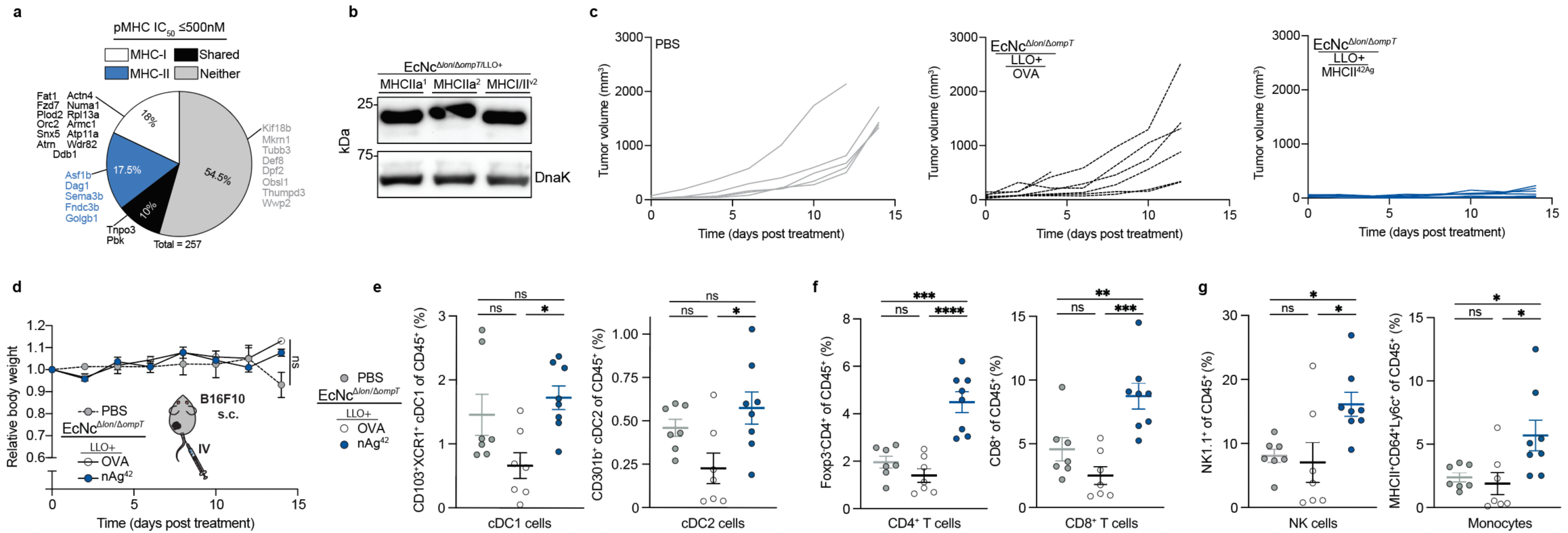
Assessment of engineered microbial neoantigen therapeutics in B16F10 melanoma. **a**, Percentage of predicted B16F10 neoantigens containing mutant-epitope(s) with ≤500nM MHC-I affinity (MHC-I), MHC-II affinity (MHC-II), both MHC-I and MHC-II affinity (Shared), or no epitope meeting affinity criteria (Neither). Previously validated neoantigens within the set are labeled. **b**, Immunoblot of B16F10 neoantigen construct expression in EcNc^Δ*lon*/Δ*ompT*/LLO+^. **c**–**g,** C57BL/6 mice with established hind-flank B16F10 melanoma tumors were treated 9 days after tumor engraftment. **c** and **d**, Every 3-5 days, mice (*n* = 5-7 per group) received intravenous injection of PBS, EcNc^Δ*lon*/Δ*ompT*/LLO+^ OVA, or the 7-strain combination EcNc^Δ*lon*/Δ*ompT*/LLO+^ nAg^42^. **c**, Individual tumor trajectories after intravenous treatment with indicated therapeutic. **d**, Relative body weight of B16F10-tumor bearing mice (*n* = 5-7 per group, ns = not significant (*P* > 0.05), two-way ANOVA with Tukey’s multiple comparisons test). **e**–**g**, On day 9 and 12 post-engraftment, B16F10 tumor-bearing mice received an intravenous injection of PBS, EcNc^Δ*lon*/Δ*ompT*/LLO+^ OVA, or the 7-strain combination EcNc^Δ*lon*/Δ*ompT*/LLO+^ nAg^42^. **g**, Left: Frequency of CD103^+^XCR1^+^ cDC1 cells in tumors (*n* = 7-8 mice per group, \**P* = 0.0132, ns = not significant (*P* > 0.05), one-way ANOVA with Tukey’s multiple comparisons test). Right: Frequency of CD310b^+^ cDC2 cells in tumors (*n* = 7-8 mice per group, \**P* = 0.0162, ns = not significant (*P* > 0.05), one-way ANOVA with Tukey’s multiple comparisons test). **h**, Left: Frequency of Foxp3^-^CD4^+^ T cells in tumors (*n* = 7-8 mice per group, \*\*\**P* = 0.0001, \*\*\*\**P* < 0.0001, ns = not significant (*P* > 0.05), one-way ANOVA with Tukey’s multiple comparisons test). Right: Frequency of CD8^+^ cytotoxic T cells in tumors (*n* = 7-8 mice per group, \*\**P* = 0.0097, \*\*\**P* = 0.0002, ns = not significant (*P* > 0.05), one-way ANOVA with Tukey’s multiple comparisons test). **i**, Left: Frequency of NK1.1^+^ NK cells in tumors (*n* = 7-8 mice per group, \**P* = 0.0189, \**P* = 0.0389, ns = not significant (*P* > 0.05), one-way ANOVA with Tukey’s multiple comparisons test). Right: Frequency of MHCII^+^CD64^+^Ly6c^+^ monocytes in tumors (*n* = 7-8 mice per group, \**P* = 0.0230, \**P* = 0.0495, ns = not significant (*P* > 0.05), one-way ANOVA with Tukey’s multiple comparisons test). **d**–**g**, Data are mean ± s.e.m.

**Extended Data Figure 9.**
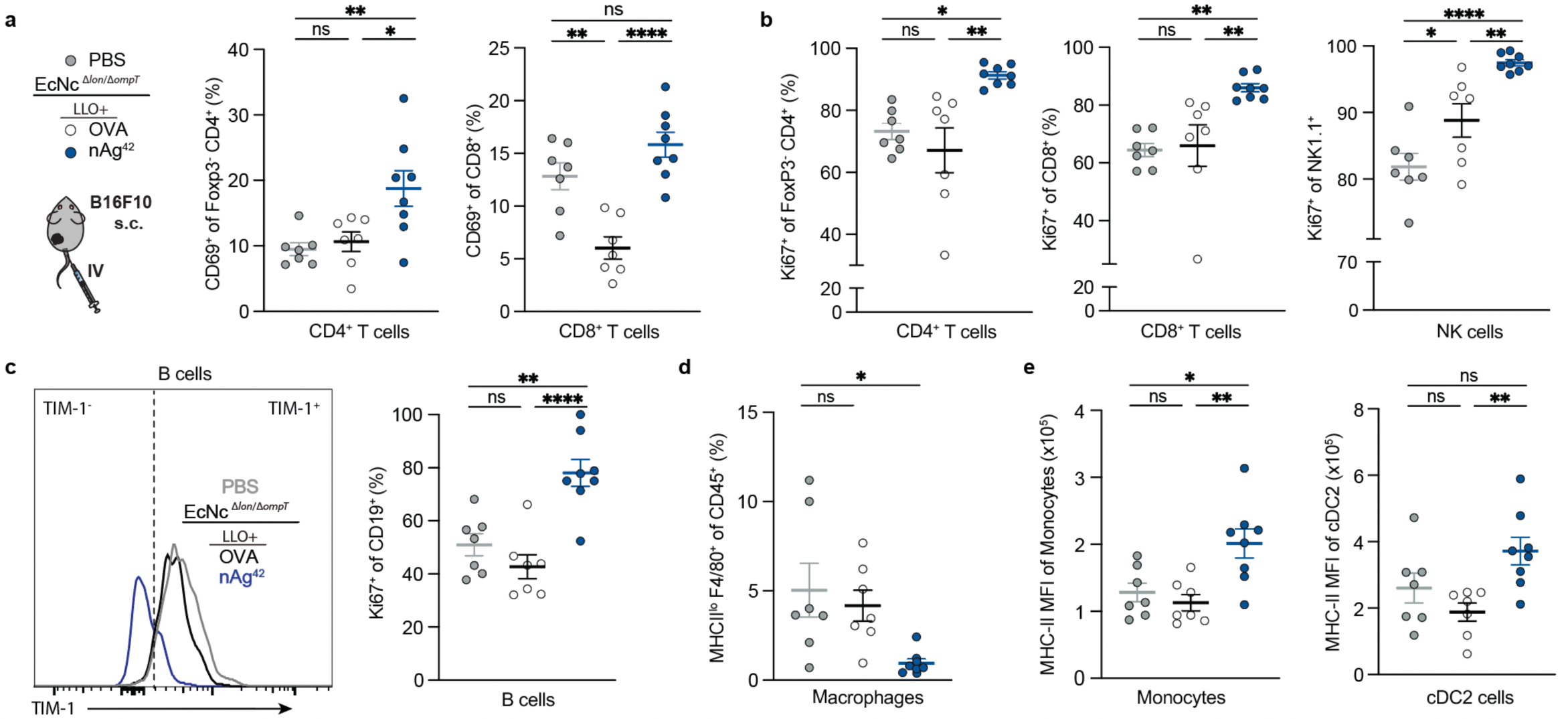
**a**–**e**, On day 9 and 12 post-engraftment, B16F10 tumor-bearing mice received an intravenous injection of PBS, EcNc^Δ*lon*/Δ*ompT*/LLO+^ OVA, or the 7-strain combination EcNc^Δ*lon*/Δ*ompT*/LLO+^ nAg^42^. **a**, Left: Percentage CD69^+^ of Foxp3^-^CD4^+^ T cells in tumors (*n* = 7-8 mice per group, \**P* = 0.0228, \*\**P* = 0.0092, ns = not significant (*P* > 0.05), one-way ANOVA with Tukey’s multiple comparisons test). Right: Percentage CD69^+^ of CD8^+^ T cells in tumors (*n* = 7-8 mice per group, \*\**P* = 0.0021, \*\*\*\**P* < 0.0001, ns = not significant (*P* > 0.05), one-way ANOVA with Tukey’s multiple comparisons test). **b**, Left: Percentage Ki-67^+^ of Foxp3^-^CD4^+^ T cells in tumors (*n* = 7-8 mice per group, \**P* = 0.0188, \*\**P* = 0.0020, ns = not significant (*P* > 0.05), one-way ANOVA with Tukey’s multiple comparisons test). Middle: Percentage Ki-67^+^ of CD8^+^ T cells in tumors (*n* = 7-8 mice per group, \*\**P* = 0.0048, \*\**P* = 0.0086, ns = not significant (*P* > 0.05), one-way ANOVA with Tukey’s multiple comparisons test). Right: Percentage Ki-67^+^ of NK1.1^+^ NK cells in tumors (*n* = 7-8 mice per group, \**P* = 0.0366, \*\**P* = 0.0070, \*\*\*\**P* < 0.0001, ns = not significant (*P* > 0.05), one-way ANOVA with Tukey’s multiple comparisons test). **c**, Left: Representative histogram of TIM-1 expression on CD19^+^ B cells in respective groups. Right: Percentage Ki-67^+^ of CD19^+^B cells in tumors (*n* = 7-8 mice per group, \*\**P* = 0.0015, \*\*\*\**P* < 0.0001, ns = not significant (*P* > 0.05), one-way ANOVA with Tukey’s multiple comparisons test). **d**, Frequency of MHC-II^lo^F4/80^+^ macrophages in tumors (*n* = 7-8 mice per group, \**P* = 0.0130, ns = not significant (*P* > 0.05), one-way ANOVA with Dunnett’s multiple comparisons test). **e**, Left: MHC-II MFI of CD64^+^Ly6c^+^ monocytes in tumors (*n* = 7-8 mice per group, \**P* = 0.0171, \*\**P* = 0.0041, ns = not significant (*P* > 0.05), one-way ANOVA with Tukey’s multiple comparisons test). Right: MHC-II MFI of CD301b^+^ cDC2 in tumors (*n* = 7-8 mice per group, \*\**P* = 0.0090, ns = not significant (*P* > 0.05), one-way ANOVA with Tukey’s multiple comparisons test). **a**–**e**, Data are mean ± s.e.m.

**Extended Data Figure 10.**
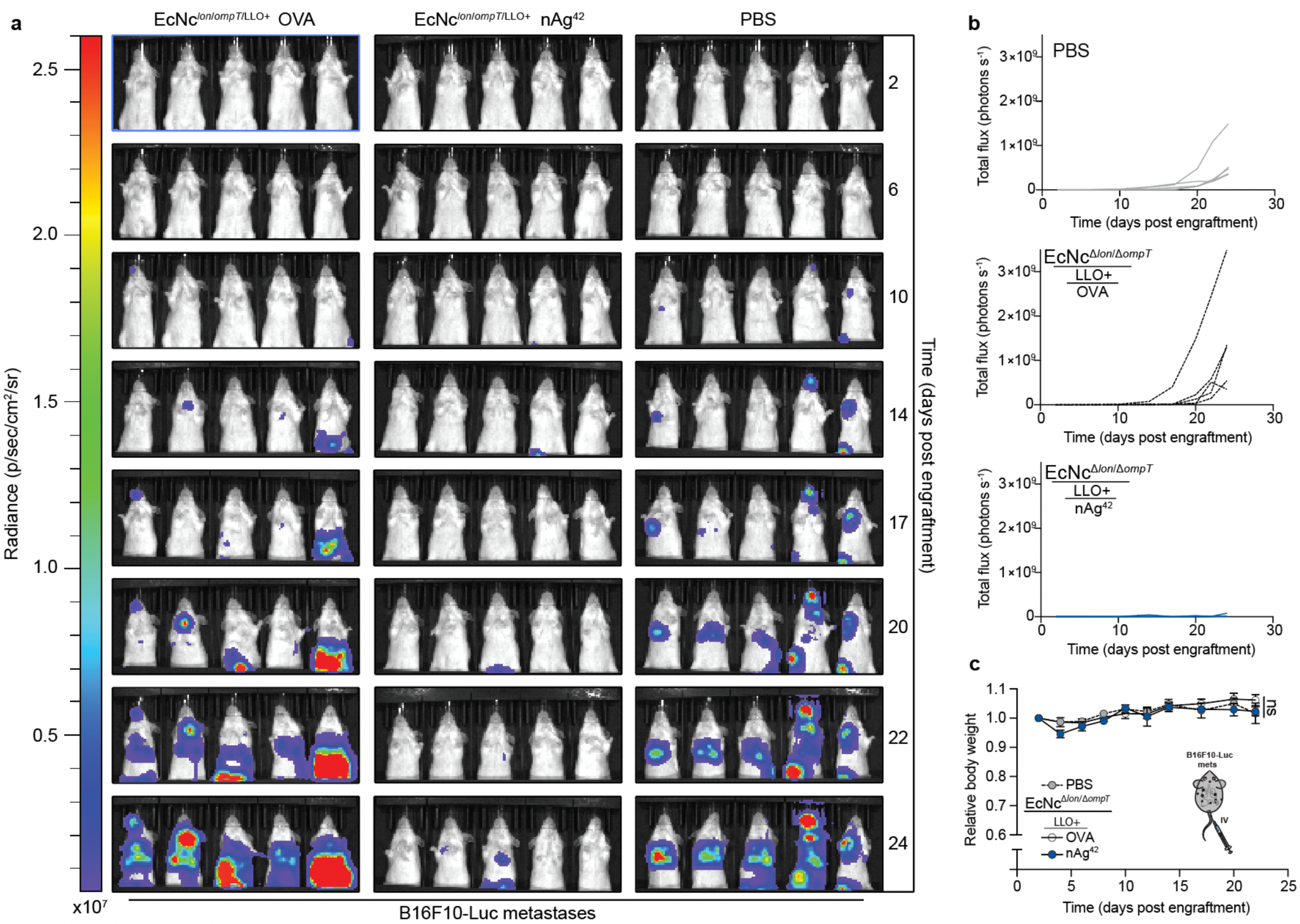
Microbial neoantigen vaccines suppress metastatic B16F10 melanoma. **a**–**c**, Mice received intravenous injection of either PBS, EcNc^Δ*lon*/Δ*ompT*/LLO+^ OVA or nAg^42^ every 3-5 days starting 2 days post intravenous injection of B16F10-Luc cells. **a**, Images of systemic metastases luminescence in each mouse in all groups over treatment course. **b**, Individual systemic metastases luminescence trajectories (*n* = 5 mice per group). **c**, Relative body weight of mice (*n* = 5 per group) after intravenous treatment with indicated therapeutic. Data are mean ± s.e.m.

**Extended Data Table 1:**
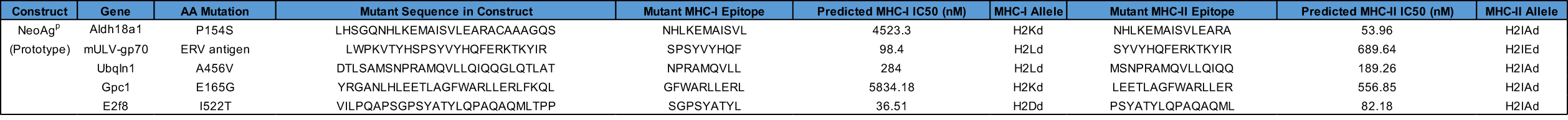
CT26 peptides in prototype neoantigen construct. Used for microbial production optimization studies.

**Extended Data Table 2:**
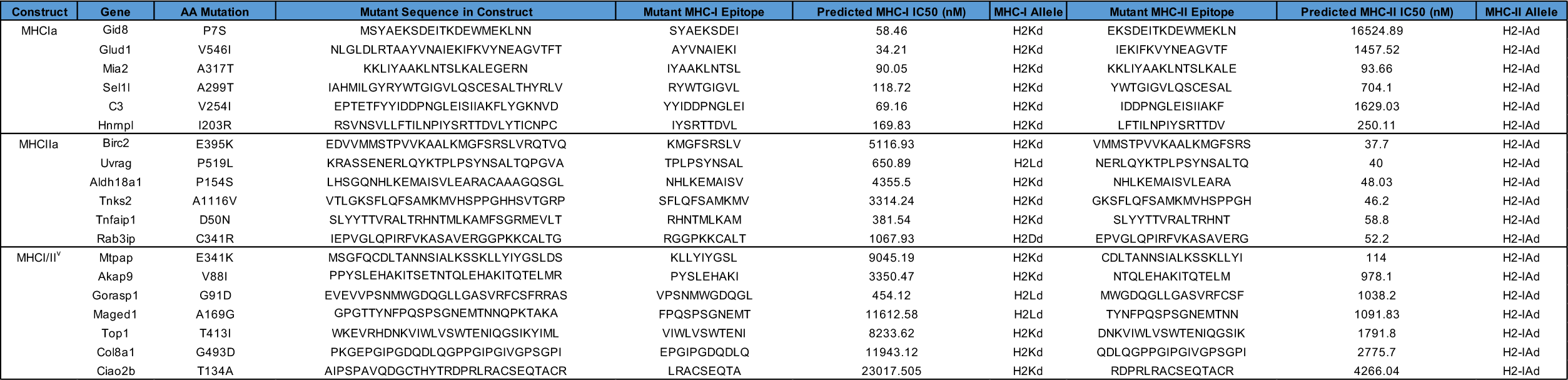
CT26 mutant peptides in neoantigen constructs. 6-7 predicted neoantigens were included in each construct. All 3 groups in combination represent nAg^19^.

**Extended Data Table 3:**
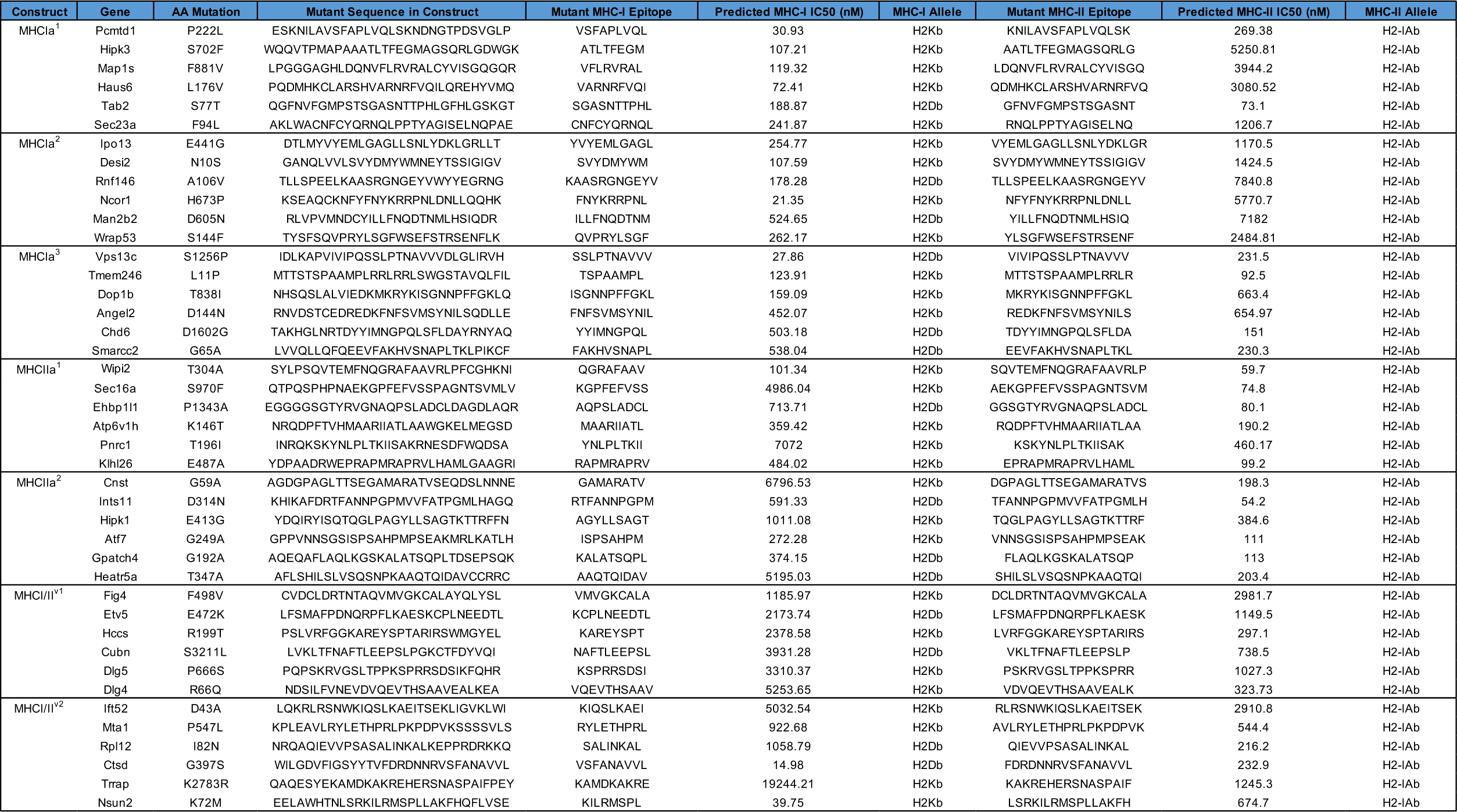
B16F10 mutant peptides in neoantigen constructs. 6 predicted neoantigens were included in each construct. All 7 groups in combination represent nAg^42^.

## Methods

### Cell lines

The B16F10 melanoma (ATCC CRL-6475) and CT26 (ATCC CRL-2638) colon carcinoma cell lines were purchased from ATCC. CT26-Luc and B16F10-Luc cells were lentivirally transduced with luciferase. Cells were confirmed mycoplasma free. Cells were cultured in incubators at 37°C with atmosphere of humidified 5% CO2. B16F10 cells were grown in DMEM supplemented with 10% (vol/vol) FBS, 1× GlutaMax, 1% (vol/vol) MEM Non-essential amino acids solution (Gibco-11140050), and 100 U/ml penicillin-streptomycin. CT26 and CT26-Luc cells were grown in RPMI-1640 supplemented with 10% (vol/vol) FBS, 1× GlutaMax, 1% (vol/vol) MEM Non-essential amino acids solution, and 100 U/ml penicillin-streptomycin.

### Exome sequencing

Paired tumor and tail DNA from BALB/c mice bearing subcutaneous CT26 tumors or C57BL/6 mice bearing subcutaneous B16F10 tumors was extracted in triplicate (*n* = 3 mice per tumor line) using Qiagen DNeasy Blood & Tissue Minikit per manufacturer’s instructions. Exome capture from mouse tumor and tail DNA triplicates was conducted using Agilent SureSelectXT All Exon kit for target enrichment DNA library preparation^65^, according to manufacturer’s instructions (Agilent). Genomic DNA was fragmented by acoustic shearing with a Covaris S220 instrument. Fragmented DNAs were cleaned, end repaired, and adenylated at the 3’-end. Adapters were ligated to DNA fragments, and adapter-ligated DNA fragments enriched with limited cycle PCR. Adapter-ligated DNA fragments were validated using Agilent TapeStation (Agilent) and quantified using Qubit 2.0 Fluorometer (ThermoFisher Scientific) and Real-Time PCR (KAPA Biosystems). Sequencing libraries were clustered onto a lane of a flow cell. After clustering, the flow cell was loaded on an Illumina HiSeq4000 Instrument per manufacturer’s instructions. Samples were sequenced using 2×150bp Paired End configuration. Image analysis and base calling was conducted by the HiSeq Control Software. Raw sequence data (.bcl files) generated from Illumina HiSeq was converted into fastq files and de-multiplexed using Illumina bcl2fastq2.17. Sequence reads were trimmed to remove adapter sequences and nucleotides with poor quality using Trimmomatic v.0.39^66^. Trimmed reads were aligned to the GRCm38 reference genome using the Illumina Dragen Bio-IT platform. Alignments were sorted and PCR/optical duplicates marked for generation of BAM files. Somatic single nucleotide variants (SNV) and insertion/deletion (indel) variants were called using Illumina Dragen^67^ and GATK Mutect2^68^. All variants from paired-normal tissue and murine variants from dbSNP^69^ were removed during the process. VCF files were left aligned and normalized, with splitting of multiallelic sites into multiple sites using bcftools 1.13^70^. Only tumor-specific variants called by both algorithms were used for further analysis.

### RNA sequencing

Tumor RNA from BALB/c mice bearing subcutaneous CT26 tumors or C57BL/6 mice bearing subcutaneous B16F10 tumors was extracted in triplicate using Qiagen RNeasy Minikit per manufacturer’s instructions. Extracted RNA samples were quantified using Qubit 2.0 Fluorometer (Life Technologies) and RNA integrity checked using Agilent TapeStation 2400 (Agilent). RNA sequencing libraries were prepared using the NEBNext Ultra RNA library Prep Kit for Illumina per manufacturer’s instructions (New England Biolabs). mRNAs were enriched with Oligo(dT) beads. Enriched mRNAs were fragmented for 15 minutes at 94°C. First strand and second strand cDNAs were synthesized subsequently. cDNA fragments were end-repaired and adenylated at 3’-ends, and universal adapters ligated to cDNA fragments, followed by index addition and library enrichment by limited-cycle PCR. Sequencing libraries were validated on Agilent TapeStation (Agilent), and quantified using Qubit 2.0 Fluorometer (Invitrogen) and quantitative PCR (KAPA biosystems). Library loading, sequencing, and read trimming were done as described above. Trimmed reads were aligned to the mm10 reference using STAR aligner v.2.5.2b^71^. Unique gene hit counts were calculated using feature Counts from Subread Package v.1.5.2. Unique reads that fell within exon regions were counted. The gene hit counts table was used for expression analysis using DESeq2^72^.

### Neoantigen prediction and selection

Mutation-specific RNA expression and allele fraction were added to somatic VCF files using Bam-readcount^73^ and VAtools (http://vatools.org). Somatic VCFs were annotated with The Ensembl Variant Effect Predictor (VEP Ensembl version 104)^74^. Only PASS variants from VCFs were considered. Annotated VCFs were analyzed using pVacSeq for neoepitope discovery^75^. MHC-I affinities were predicted with NetMHCpan version 4.1^76^ and NetMHC version 4.1^77^, and MHC-II affinities predicted with NetMHCIIpan version 4.1^78^ and NNalign version 2.0^79^. Exonic mutations based on single-nucleotide polymorphisms (SNP) or indels predicted to generate mutant MHC-binding peptides were included based on the set of minimum criteria: 1) present in all tumor sample triplicates (DNA-VAF ≥0.05) and none of the normal tissue triplicates, 2) non-synonymous mutation resulting from either SNP or indel, 3) confirmed exonic mutation transcription (RNA-VAF ≥0.05) and gene expression by RNA-sequencing in tumor sample triplicate (TPM ≥1), 4) at least 1 epitope of predicted MHC-I or MHC-II IC50 ≤6000nM. Predicted neoantigens fulfilling all prior constraints and which satisfied the following calculation were then considered for selection and incorporation into therapeutics:

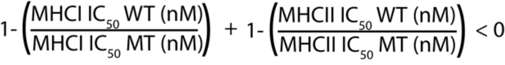

Where MHC-I IC50 MT and WT are the predicted MHC-I IC50 of the highest affinity MHC-I binding mutant and corresponding wildtype epitope, and MHC-II IC50 MT and WT are the predicted MHC-II IC50 of the highest affinity MHC-II binding mutant and corresponding wildtype epitope contained within the exonic mutation-derived long peptide.

## Strains and Plasmids

Plasmids were constructed using restriction-enzyme mediated and Gibson assembly cloning methods. Neoantigen construct iterations were designed and created as Geneblocks (IDT) encoding a constitutive promoter and 5’-UTR containing selected ribosome-binding site, followed by coding region comprised of mutant-residue containing LPs connected in tandem or by various linkers as indicated. 5’-*Bam*HI and 3’-*Xba*I restriction endonuclease sites were added to constructs. Coding sequences were codon optimized for *E. coli*. Constructs were cloned between *Bam*HI and *Xba*I restriction sites on a stabilized p246-*lux*CDABE plasmid where *lux*CDABE had been cloned out^80^, and flanked by 3’-λ-phage transcription terminator, with high-copy pUC origin. For protein expression assessment studies, the codon sequence for a 6x-Histidine Tag (HisTag) was added immediately before the stop-codon within the neoantigen construct coding sequence by PCR amplification of full construct plasmids with oligonucleotide containing 6x-HisTag sequence followed by kinase, ligase, *Dpn*I enzyme mix protocol (NEB). Neoantigen construct plasmids were transformed into chemically competent *E. coli* DH5α or BL21(DE3) (New England Biolabs), or electrocompetent EcN parental strain or genetic derivatives. The parental EcN strain and derivatives used in this study harbor an integrated luciferase cassette within the genome, which contains an erythromycin resistance casette^81^. Plasmid encoding constitutive Listeriolysin-O (LLO) was constructed by cloning in hok/sok stabilization system to pCG02-p15a backbone^3^, PCR amplification of backbone with SLC cloned out, and Gibson assembly of Geneblock encoding LLO under constitutive promoter and backbone. Constitutive LLO plasmids were transformed into electrocompetent EcN parental and genetic derivative strains. Strains were cultured in lysogeny broth (LB) medium with antibiotics for plasmid retention (pUC:kanamycin 50 μg/ml, p15a:spectinomycin 50 μg/ml) in a 37°C orbital incubator.

### Construction of cryptic plasmid cured EcN

EcN cryptic plasmids were cured with Cas9-mediated double-strand break, as described previously^42^. Briefly, EcN was transformed with pFREE or pCryptDel4.8 to cure the cryptic plasmids pMUT1 or pMUT2, respectively. The transformants were grown overnight and diluted 1:1000 the next day into fresh LB containing 0.2% rhamnose and 0.43μM anhydrotetracycline. After 24 hours of incubation, the culture was streaked onto LB plates without antibiotics and incubated overnight in a 30°C incubator. Colonies were screened with colony PCR to verify loss of cryptic plasmids.

### Construction of genetic knockout strains

Genetic knockouts were performed using the lambda red recombination system^82^. In brief, EcNc was transformed with pKD46. Transformants were grown at 30°C in LB with ampicillin and L-arabinose, then made electrocompetent. The chloramphenicol resistance cassette with corresponding overhangs for each target gene for deletion was prepared by PCR amplification of pKD3. Electroporation was performed using 100μl of competent cells and 50-300ng amplified DNA. After 2 hours of recovery, cells were plated on LB agar containing chloramphenicol and incubated at 37°C overnight. Target gene deletion was verified by colony PCR. For excision of the antibiotic resistance marker, pCP20 was transformed, transformants plated on fresh LB plates containing ampicillin and incubated at 30°C overnight. Selected colonies were then inoculated onto fresh LB plates without antibiotics and cultured at 43°C overnight for induction of flippase and plasmid curing. Clones were subsequently screened for loss of antibiotic resistance.

### qPCR for plasmid copy number

Copy number variant plasmids were constructed from a high-copy pUC-GFP^80^ plasmid. The plasmid backbone excluding the pUC origin was PCR-amplified and Gibson assembled with sc101*, p15A, or ColE1 origin of replication insert. The respective inserts were prepared from PCR amplification of template plasmid pCG02_sc101*, pCG02_p15A, or pTH05_ColE1. Plasmid copy number (PCN) was determined as reported previously^80^, where relative abundance of plasmid DNA compared to genomic DNA is measured by qPCR. Briefly, strains with the plasmid of interest were grown at 37°C overnight in fresh LB with appropriate antibiotics. Cells were harvested by centrifugation at 3,000rcf at 4°C for 10 minutes, supernatant removed and cell pellet resuspended in distilled water to optical density measurement at 600nm (OD600) = 1. Resuspended cells were 5-fold serially diluted. Samples were denatured at 95°C for 10 minutes and 2μl of each sample dilution was added into 18μl of NEB Luna Universal qPCR Master Mix in each well of a 96-well plate. 25-fold diluted samples were used for the measurement of crossing point (Ct) values: the cycle number when amplified sample fluorescence exceeds background. 5-fold diluted samples were used for generation of the standard curve for PCR efficiency (E). E was defined from the slope (S) of each standard curve with the equation E = 5(-1/S), and plasmid copy number (PCN) was determined with the equation:

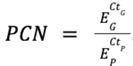

Where respective values for genomic DNA are denoted by subscript G and plasmid DNA by subscript P.

### Immunoblot and ELISA

Strains expressing neoantigen construct with C-terminal 6x-HisTag were grown overnight in LB media with appropriate antibiotics. Equalization of OD600 measurement to match CFU/ml between all cultures was done prior to all sample processing. CFU-matched cultures were centrifuged at 3000rcf at 4°C for 10 minutes. For immunoblot, samples were resuspended in B-PER lysis reagent (ThermoFisher Scientific) containing 250U/ml benzonase nuclease (Millipore Sigma), and 1U/ml rLysozyme (Millipore Sigma) and placed on an orbital shaker for 15 minutes at room temperature. Samples were centrifuged at 10,000rcf for 20 minutes at 4°C to separate soluble and insoluble fractions, or total lysate used directly. Processed samples were mixed with SDS-loading buffer with 5mM dithiothreitol, boiled, and subject to immunoblot analysis. For relative quantification of immunoblot chemiluminescent intensity, target proteins bands on the same blot were normalized to the loading control band DnaK for the same sample. Normalized values were divided to provide relative intensity values. Mouse anti-6xHis (αTHE) was purchased from Genscript, mouse anti-DnaK was purchased from Abcam (8E2/2).

For HisTag ELISA, samples were resuspended in ice-cold PBS containing HALT^TM^ protease inhibitor cocktail (ThermoFisher Scientific). Samples were sonicated on ice for 2 minutes total time. Sonicated samples were centrifuged at 10,000rcf for 20 minutes at 4°C. Soluble sample fractions were analyzed using GenScript HisTag ELISA Detection Kit per manufacturer’s instructions.

For *ex vivo* immunoblot analysis, BALB/c mice bearing established hind-flank CT26 tumor were injected intravenously with EcNc*^Δlon/ΔompT^*^/LLO+^ nAg^19-His^ strain cocktail where each construct (MHCIa, MHCIIa, and MHCI/II^v^) contained a C-terminal 6x-HisTag. 48 hours after treatment, tumors and TDLNs were extracted from 5 mice and placed in B-PER lysis reagent (ThermoFisher Scientific) with 250U/ml benzonase nuclease (Millipore Sigma), and homogenized using a gentleMACS tissue dissociator (Miltenyi Biotec, C-tubes). Tissue homogenate was sonicated on ice for 3 minutes. Sonicated samples were centrifuged at 10,000rcf for 20 minutes at 4°C to separate soluble and insoluble fractions and diluted in lysis buffer. Sample fractions were mixed with SDS-loading buffer with 5mM dithiothreitol, boiled, and subject to immunoblot analysis.

### Blood bactericidal assay

EcN wildtype or EcNc^Δ*lon*/Δ*ompT*^ were cultured overnight in LB media without antibiotics. Cultures were centrifuged at 3000rcf for 10 minutes, resuspended in 1ml of ice-cold sterile PBS, and normalized to OD600 = 1. 50 μl of OD600 = 1 microbe suspension was added to 1ml of single donor human whole blood (Innovative Research) in triplicate and incubated in a 37°C stationary incubator. After 2 hours incubation, a sample was taken from each blood-microbe mixture and serial dilution prepared in PBS. Dilutions were plated on LB agar with erythromycin (25 μg/ml). After incubation overnight at 37°C, colonies were quantified by spot-forming assay and CFU/ml blood calculated.

### Biofilm assay

Biofilm formation assays were conducted as described previously^83^. Briefly, EcN wildtype, cryptic plasmid cured (EcNc), Lon knockout (EcNc*^Δlo^*^n^), OmpT knockout (EcNc*^ΔompT^*) or double protease knockout (EcNc*^Δlon/ΔompT^*) were cultured 48 hours in LB media with 25 μg/ml erythromycin in borosilicate glass tubes in a 30°C stationary incubator, with tube caps wrapped with parafilm to prevent evaporation. At 48 hours, cultures were discarded and borosilicate tubes washed 3-times with PBS. Tubes were inverted and let dry for 6-hours. Biofilms left on borosilicate tubes were stained with 0.1% (vol/vol) crystal violet for 15 minutes. Crystal violet stain was discarded, tubes washed 3-times with PBS, then inverted and let dry overnight. Crystal violet-stained biofilms were dissolved with 95% ethanol and transferred to 96-well plates for measurement of absorbance at 590nm.

### Phagocytosis assay

Bacterial phagocytosis assays were adapted from previous work^84^. Culture and isolation of murine bone marrow derived macrophages (BMDMs) was performed as described previously^85^. Bulk femoral bone marrow cells from BALB/c or C57BL/6 mice were cultured on 15-cm non-tissue culture treated Petri dishes in RPMI with 20% FBS, 25ng/ml M-CSF (R&D Systems), and 100 U/ml penicillin-streptomycin. Media was replaced with fresh media after 4 days of culture. After 7 days in culture, plates were washed with PBS and adherent macrophages were dissociated using trypsin-EDTA. Macrophages were washed in PBS, resuspended at a density of 2×10^5^/ml in media, and 1ml transferred to each well of 24-well plates. 24-well plates were incubated overnight in a 37°C incubator with humidified 5% CO2. EcN wildtype or EcNc*^Δlon/ΔompT^*, with or without a constitutive GFP-expressing plasmid were cultured overnight in LB media with appropriate antibiotics. Bacterial cultures were centrifuged at 3000rcf for 10 minutes, washed 3-times with sterile PBS, and resuspended at a density of 4×10^8^ bacteria/ml in sterile PBS. Media from wells containing adherent macrophages was aspirated, wells washed 3-times with PBS, and 1ml of RPMI with 5% mouse serum added to each well. Latrunculin A was added at a concentration of 1μM to selected wells to inhibit phagocytosis. 2×10^7^ CFU of microbes were added to each well with each condition tested in triplicate. Microbial strains were incubated with BMDMs for 30-minutes in a 37°C incubator at 20rpm. After 30 minutes, media was aspirated and wells washed 6-times with sterile ice-cold PBS. Adherent macrophages were dissociated using non-enzymatic cell dissociation buffer (Gibco), resuspended in FACS buffer (PBS containing 2% FBS, 2mM EDTA, and 0.09% sodium azide) and analyzed by flow cytometry.

### *In vitro* BMDM activation

BMDMs were cultured as described above for phagocytosis assays. BMDMs were washed in PBS, resuspended at a density of 2×10^5^/ml in media, and 1ml transferred to each well of 24-well plates. 24-well plates were incubated overnight in a 37°C incubator with humidified 5% CO2. Wildtype EcN or EcNc*^Δlon/ΔompT^* with constitutive expression of OVA from a pUC origin plasmid were cultured overnight in LB media with appropriate antibiotics. Cultures were centrifuged at 3000rcf for 10 minutes, washed 3-times with PBS, and resuspended at a density of 4×10^8^ bacteria/ml in sterile PBS. Media from wells containing macrophages was aspirated, wells washed 3-times with PBS, and 1ml of RPMI with 5% mouse serum added to each well. 1×10^7^ live microbes were added to each well, with each condition replicated in triplicate. Live microbial strains were incubated with BMDMs for 6-hours in a 37°C incubator. After 6-hours, media was aspirated and wells washed 6-times with sterile ice-cold PBS. Adherent macrophages were dissociated using non-enzymatic cell dissociation buffer (Gibco), resuspended in FACS buffer, and analyzed by flow cytometry. DRAQ7 cell viability reagent was used to exclude dead cells (diluted 1:1000 in FACS buffer). Extracellular antibodies for BMDM activation panel included CD80 (16-10A1, Biolegend), MHC-II (M5/114.15.2, Biolegend), PD-L1 (10F.9G2, Biolegend), and H2K^b^-SIINFEKL (25-D1.16, Biolegend).

### *In vitro* BMDC stimulation

BMDC isolation and culture from mouse bone marrow was adapted from previous methods^86^. BMDC from C57BL/6 mice were cultured on 15-cm non-tissue culture treated Petri dishes in RPMI with 20% FBS, 20ng/ml GM-CSF (Biolegend), and 100 U/ml penicillin-streptomycin. Every 1-2 days for the first 4 days, plates were gently washed and non-adherent granulocytes removed by aspirating 50% of the culture media with subsequent replacement of fresh media. On day 4, media was aspirated completely and replaced with fresh culture media with 20ng/ml GM-CSF. On day 6, BMDC plates were washed with PBS and loosely adherent and non-adherent cells harvested. Cells were centrifuged at 300rcf for 5 minutes, resuspended in fresh culture media, and replated on 15-cm non-tissue culture treated Petri dishes. On day 7-8, plates were washed with PBS and loosely adherent and non-adherent cells harvested. Cells were centrifuged at 300rcf × 5 minutes, resuspended in fresh culture media at a density of 2.5×10^5^/ml, and 200μl transferred to 96-well plates and incubated overnight in a 37°C incubator. The next day, media from wells containing BMDCs was aspirated, and 1ml of RPMI with 5% mouse serum was added to each well. BMDCs were pulsed with live bacteria at an MOI of 10 for 2-hours. Plates were centrifuged at 300rcf for 5 minutes, media aspirated and replaced with fresh RPMI with 10% FBS, 10μg/ml gentamicin and 100 U/ml penicillin-streptomycin. Gentamicin concentration was increased to 40μg/ml after 2-4 hours. Plates were incubated for 12-48 hours in a 37°C incubator, after which time supernatant was assessed for IL-12p70 using the Mouse IL-12p70 Quantikine ELISA Kit (R&D systems) according to the manufacturer’s instructions.

### OT-I and OT-II T cell stimulation and proliferation

BMDC were cultured as above, resuspended at a density of 2.5×10^5^/ml, and 5×10^4^ BMDC transferred to 96-well plates and incubated overnight in a 37°C incubator. The next day, media from wells containing BMDCs was aspirated, and 1ml of RPMI with 5% mouse serum was added to each well. BMDCs were pulsed with 2×10^6^ CFU of the respective bacterial strain for 2.5-hours, plates were centrifuged at 300rcf × 5 minutes, media aspirated and replaced with fresh RPMI with 10% FBS and 10μg/ml gentamicin and 100 U/ml penicillin-streptomycin. Gentamicin concentration was increased to 40μg/ml after 2-4 hours. Spleens from naïve OT-I and OT-II mice were extracted, filtered through 100µm cell strainers and washed in complete RPMI (RPMI-1640 supplemented with 10% (vol/vol) FBS, 1× GlutaMax, 1% (vol/vol) MEM Non-essential amino acids solution (Gibco-11140050), and 100 U/ml penicillin-streptomycin). OT-I and OT-II T cells were isolated from single-cell suspensions of spleens from the respective transgenic mouse using the EasySep™ Mouse T Cell Isolation Kit (StemCell Technologies) according to the manufacturer’s instructions. Purified OT-I and OT-II T cells were resuspended in T cell media (complete RPMI supplemented with 50μM β-mercaptoethanol) at a density of 5×10^5^/ml, and 5×10^4^ T cells incubated with 5×10^4^ BMDC pulsed with the respective microbial strains. For cytokine secretion assessment, T cells were incubated with BMDCs for 24 hours, after which time supernatant was assessed for IFN-γ and IL-2 using Mouse IFNgamma Quantikine ELISA Kit and Mouse IL-2 Quantikine ELISA Kit (R&D systems) according to the manufacturer’s instructions.

CFSE proliferation assays were conducted as previously described^87^. 1×10^7^ OT-I or OT-II T cells were resuspended in 1ml of room temperature PBS, and 1μl of 5mM CFSE (Biolegend) was added. T cells were incubated in CFSE solution for 5 minutes at room-temperature protected from light, after which time staining was quenched by adding 10-times the staining volume of cell culture media. T cells were centrifuged at 300rcf for 5 minutes, resuspended in T cell media at a density of 5×10^5^/ml and incubated for an additional 10 minutes at room temperature. 5×10^4^ T cells were incubated with 5×10^4^ BMDC pulsed with the respective live microbial strains. At 48 hours, 50% of media from each well was gently aspirated and replaced with fresh T cell media. At 72-96 hours, OT-I and OT-II T cells were harvested and analyzed via flow cytometry. DRAQ7 cell viability reagent was used to exclude dead cells (diluted 1:1000 in FACS buffer). Extracellular antibodies for CFSE assays included CD3 (17A2, Biolegend), CD8a (53-6.7, Biolegend), and CD4 (GK1.5, Biolegend).

### Listeriolysin hemolytic activity assay

Sheep red blood cell (RBC) lysis by bacterial lysate was performed as described previously^88^. Briefly, bacteria were grown overnight in fresh LB containing appropriate antibiotics. Cultures were centrifuged at 3000rcf for 10 minutes, supernatants discarded, and cell pellet resuspended to OD600 = 8 in 0.1% (w/w) BSA in sterile PBS titrated to pH of 5.25 with 1M HCl. Bacteria were sonicated for 2 minutes. After sonication the soluble fraction was isolated by centrifugation at 10000rcf at 4°C for 20 minutes. Sheep RBCs were washed 3-times with PBS and resuspended at a final concentration of 6×10^8^/ml in 0.1% (w/w) BSA in PBS titrated to pH of 5.25. Equal parts of bacterial lysate soluble fraction and sheep RBC suspension were mixed and incubated for 15 minutes at 37°C. After incubation RBC mixtures were centrifuged at 1000rcf for 1 minute at 4°C and supernatant absorbance at 541nm measured to quantify RBC lysis.

### Animal experiments

All animal experiments were approved by the Institutional Animal Care and Use Committee (Columbia University, protocol AABQ5551). Female 6–7-week-old BALB/c, C57BL/6, and B6(Cg)-Tyrc-2J/J (Jackson labs) mice were kept in accordance with rules for animal research at Columbia University. For subcutaneous tumor models: 5×10^6^ CT26 cells in 100μl sterile PBS were inoculated subcutaneously on the hind-flank of BALB/c mice, or 5×10^5^ B16F10 melanoma cells in 100μl sterile PBS subcutaneously on the hind flank (orthotopic) of C57BL/6 mice using a 26G-needle on a 1cc syringe. CT26 tumors were allowed to establish as indicated for each experiment, and mice distributed between groups to equate average starting tumor volume before treatment. B16F10 orthotopic tumors were allowed to establish for 10 days, and initial tumor volume equated between groups before treatment. Tumor dimensions were measured unblinded with a caliper every 1-3 days for calculating tumor volumes using the equation (*a*^2^ *x b*)/*2* (*a* = width, *b* = length, where width is the smaller dimension). Group tumor sizes were computed as mean ± s.e.m. Body weight was measured each time tumor measurements were taken. Animals were euthanized when either: tumor burden >2 cm in largest dimension for any subcutaneous tumor, >20% body weight loss, as otherwise recommended by veterinary staff or exhibiting clinical signs of impaired health.

For systemic metastases models, 5×10^5^ CT26-Luc cells or 1.5×10^5^ B16F10-Luc cells were injected in 100μl sterile PBS through the lateral tail vein with a 27G-needle on 1cc syringe. Metastases were allowed to establish for 4 days in Balb/C mice prior to treatment for CT26-Luc, and for 2 days in C57BL/6 albino mice (B6(Cg)-Tyrc-2J/J) for B16F10-Luc. Mice were randomly distributed between groups after metastases engraftment and prior to treatment. For *in vivo* luminescence tracking of metastases burden, mice were injected i.p. with 125μl aqueous solution of D-Luciferin (50mg/ml) 6 minutes prior to imaging, and placed under isoflurane anesthesia for imaging using an *in vivo* imaging system (IVIS). Total flux from the lungs (CT26-Luc) or body (B16F10-Luc) was used to quantify tumor burden.

### Microbial administration for *in vivo* experiments

For therapeutic administration, bacterial strains were grown overnight in fresh LB media containing the appropriate antibiotics. Overnight cultures were centrifuged at 3000rcf at 4°C for 10 minutes and washed 3-times with ice-cold sterile PBS. Microbes were delivered intratumorally at a concentration of 5 × 10^8^ CFU/ml in sterile PBS, with 20μl injected using a 1cc syringe with 29G-needle. For intravenous treatment, 100μl of microbes were delivered at a concentration of 1 × 10^8^ CFU/ml in sterile PBS, through the lateral tail vein using a 1cc syringe with 29G-needle.

### Biodistribution and *in vivo* bacterial dynamics

For biodistribution experiments, BALB/c mice bearing established hind-flank CT26 or lung metastasis CT26-Luc tumors were injected intravenously with 100μl of 1 × 10^8^ CFU/ml EcNc*^Δlon/ΔompT^*^/LLO+^. 96-120 hours after single IV injection for hind-flank tumors or at sac-point for lung metastases, tumors or tumor-bearing lungs and organs were extracted from mice, weighed, and homogenized using a gentleMACS tissue dissociator (Miltenyi Biotec, C-tubes). Homogenates were serially diluted in sterile PBS and plated on LB agar plates at 37°C overnight. Colonies were quantified per organ and computed as CFU per gram of tissue. For tracking bacterial colonization of subcutaneous tumors via microbial luminescence, tumor-bearing mice treated intratumorally or intravenously with wildtype EcN parental strain or genetic derivates were imaged using IVIS at various time points. For abscopal experiments, treated and untreated tumors were harvested 14 days after a single intratumoral bacterial injection.

### *Ex vivo* T cell killing assay

Naïve tumor-free C57BL/6 mice were injected intravenously every 4 days with either PBS, EcNc^Δ*lon*/Δ*ompT*/LLO+^ OVA or nAg^42^ for a total of 4 doses. 5 days after the final dose spleens from treated mice were extracted, filtered through 100µm cell strainers, and washed in complete RPMI. T cells were isolated from single-cell suspensions of spleens from the respective mouse using the EasySep™ Mouse T Cell Isolation Kit (StemCell Technologies) according to the manufacturer’s instructions. Purified T cells were resuspended in T cell media for use in the specific lysis assay.

The luciferase based killing assay was adapted from previous methods^89^. B16F10-Luc target cells were grown for 48 hours in the presence of 100U/ml murine IFN-γ. Target cells were harvested and plated at 1×10^4^ cells per well in a 96 well plate. 12 hours later T cells were added to each well to achieve designated effector-to-target ratios (10:1, 20:1, or 40:1). After 42 hours co-incubation, 50U/ml murine IL-2 was added to all wells. Luminescence from each well was quantified after addition of One-Glo Luciferase Assay System (Promega), per manufacturer’s instructions, after 48-96 hours of co-culture. Minimum lysis wells contained only B16F10-Luc target cells. In maximum lysis wells, 10μL of media was replaced with 10μL of 3% Triton X-100 30 minutes before luminescence readout. Specific lysis (%) was calculated using the luminescence values of the respective conditions in the formula:

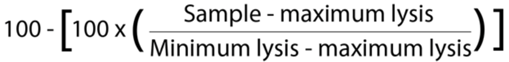

### Flow cytometry immunophenotyping

Tumors and TDLN were extracted 8 days post intravenous treatment for flow cytometry immunophenotyping. Lymphoid and myeloid immune subsets were isolated from tumor tissue by mechanical homogenization of tumor or TDLN tissue, followed by digestion with collagenase A (1mg/ml, Roche) and DNase I (0.5 µg/ml, Roche) in isolation buffer (RPMI 1640 with 5% FBS, 1% L-glutamine, 1% penicillin-streptomycin, and 10mM HEPES) for 1 hour at 37°C for tumors or 30 minutes at 37°C for TDLNs, on a shaker platform at 150rpm. For *ex vivo* lymphocyte stimulation with PMA and ionomycin, TDLNs were not digested prior. Tumor and TDLN homogenates were filtered through 100 µm cell strainers and washed in isolation buffer. To measure cytokine production by T cells, cells were stimulated for 3-hours with PMA (50ng/ml, Sigma-Aldrich) and ionomycin (1nM, Calbiochem) in the presence of brefeldin A (1μg/ml). To measure neoantigen-specific cytokine production by T cells, cells were stimulated for 5 hours with pools of peptides (2μg/ml) representing the neoantigens encoded in therapeutic strains in the presence of brefeldin A (1μg/ml). Cells were stained in FACS buffer. Ghost Dye cell viability reagent was used to exclude dead cells (diluted 1:1000 in PBS). Extracellular antibodies for lymphoid immunophenotyping included: CD4 (RM4-5, Biolegend), NKp46 (29A1.4, BD Biosciences), NK1.1 (PK136, Biolegend), CD45 (30-F11, BD Biosciences), B220 (RA3-6B2, BD Biosciences), CD19 (6D5, Biolegend), CD8a (53-6.7, Tonbo), TIM-1 (RMT1-4, BD Biosciences), CD69 (H1.2F3, BD Biosciences). After extracellular staining, cells were washed with FACS buffer, and fixed using the FOXP3/transcription factor staining buffer set (Tonbo), as per manufacturer’s instructions. Intracellular antibodies for lymphoid immunophenotyping included: Foxp3 (FJK-16s, eBioscience), CD3ε (145-2C11, Tonbo), TCRβ (H57-507, BD Biosciences), Ki67 (SolA15, eBioscience), Granzyme-B (QA16A02, Biolegend), TNF-α (MP6-XT22, eBioscience), and IFN-γ (XMG1.2, Tonbo). For myeloid immunophenotyping, extracellular antibodies included: Ly6C (HK1.4, Biolegend), I-A/I-E (M5/114.15.2, Biolegend), XCR1 (ZET, Biolegend), CD11b (M1/70, Biolegend), CD103 (2E7, Biolegend), CD45 (30-F11, BD Biosciences), F4/80 (BM8, Biolegend), CD11c (HL3, BD Biosciences), CD172a/SIRPα (P84, Biolegend), Ly6G (1A8, Biolegend), and PD-L1 (10F.9G2, Biolegend), CD301b (URA-1, Biolegend), CD3 (145-2C11, Biolegend), CD19 (1D3, Biolegend), NK1.1 (PK136, Biolegend), CD64 (X54-5/7.1, Biolegend). After staining, cells were washed and resuspended with FACS buffer for flow cytometry analysis using a BD LSRFortessa or Cytek Aurora cell analyzer. FACS Diva or SpectroFlo software was used for data acquisition. Collected flow cytometry data were analyzed using Flowjo.

### Synthetic Peptides

Synthetic peptides representing neoantigens for lymphocyte restimulation assays were synthesized by and purchased from Peptide 2.0. All peptides were ≥95% purity.

### Statistics

Statistical analyses were performed using GraphPad Prism 9. For each experiment, the particular statistical analysis is detailed in the respective figure legend. Unpaired student’s T-test, one-way ANOVA, or two-way ANOVA with appropriate post-hoc test was used for data that were approximately normally distributed. For analysis of Kaplan-Meier survival experiments, log-rank (Mantel-Cox) test was used. All analyses were two-tailed.

## Author Contributions

A.R., T.D., and N.A. conceived the study. A.R. designed the system, conducted sequencing analysis and neoantigen selection, constructed therapeutics, and performed most experiments. A.R. and J.I. designed bacterial genetic knockouts. A.R., J.I., S.H., M.K., U.J., W.S., and A.V. performed cloning. J.I., S.H., M.K., and U.J. performed knockouts. A.R. and B.R. performed *in vitro* T cell assays. J.I., Z.S., S.H., M.K., and U.J performed blood bactericidal assays. J.I. and U.J. performed qPCR. J.I., Z.S., W.S, C.R.G., S.H., J.H., E.R.B., R.L.V., M.K. and A.V. helped with *in vivo* and/or *ex vivo* experiments. A.R. and B.R. performed immunophenotyping, F.L., M.R., and J.I. assisted. A.R., T.D., and N.A. analyzed data and wrote the manuscript with input from all authors.

## Acknowledgements

Research reported in this publication was performed in the Columbia University Department of Microbiology and Immunology Flow Cytometry core facility and the Irving Cancer Research Institute’s Flow Cytometry core facility. We thank L. Kreindler, T.M. Savage, and N. Chen for critical comments and review.

